# Empirical models of initial attack success are not suitable for triaging ignitions in Victoria, Australia

**DOI:** 10.1101/2025.07.31.668020

**Authors:** Kristy Butler, Elena Tartaglia, Jason Rennie, Stephen Deutsch, Nick McCarthy

## Abstract

**Background:** Initial attack refers to the first firefighting actions undertaken at a newly reported fire, with the intention to control the fire’s spread. When multiple ignitions are reported and resources constrained, decisions need to be made about triaging fires for an escalated response.

**Aims:** We developed empirical initial attack models for grass and forest fires for the specific purpose of triaging fires, and evaluated the models relative to their intended application.

**Methods:** We used a recently developed dataset of spreading fires and tested three statistical modelling techniques to model the probability of unsuccessful initial attack. Model performance was evaluated with case studies, including major fire episodes.

**Key results:** The models showed limited skill overall. The best performance was observed for lightning-ignited forest fires in remote areas, but performance was not adequate to support implementation on days with extreme fire behaviour.

**Conclusions:** Our approach did not yield suitable models for operational triage of fires, because there are many on-the-ground factors that influence the success of initial attack, and they are difficult to capture as data.

**Implications:** This study shows the importance of developing and testing empirical models relative to their intended application. It also demonstrates the significance of unmeasured bottlenecks in wildfire suppression on initial attack outcome.

**Summary text for Table of Contents:** This study develops models to aid operational staff in triaging fires for an escalated response. We determined that model performance was not suitable for this use case. We found that using both modelling visualisations and case studies were fundamental in assessing the model’s applicability for operations.

## Introduction

Initial attack refers to the set of firefighting actions by the first resources sent to a newly reported fire. Fire management agencies aim to establish control of fires during initial attack to prevent impacts to nearby communities, industries and the environment. When it becomes apparent that a fire is unable to be controlled in initial attack, fire management agencies undertake a series of additional actions, such as dispatching further firefighting units and aircraft, issuing and escalating community warnings and planning extended strategies and tactics. There are benefits to initiating this increased response early (Arienti *et al*. 2006), but it is inefficient to over-resource fires where initial attack actions will be successful. Sending the right number of resources becomes particularly significant on days when multiple new ignitions or existing fires compete for the same pool of resources. On these days, tools that indicate which ignitions are less likely to be controlled in initial attack can help operations staff to rapidly triage fires for an increased response.

In Victoria, Australia, the decisions to increase response beyond the default level are informed by a combination of sources. Situation reports provided by the first arriving crews to the fireground give an indication of whether the fire is likely to be controlled with the resources enroute or on the ground. However, these reports are delayed by the time taken for resources to respond and travel to the fireground, and rely on a subjective assessment by the crews. Prior to a fire starting, Victorian fire agencies use fire danger indices to inform a range of decisions around resource readiness, including standing up resources in fire stations and incident control centres and automatically dispatching aircraft to fires. While there are no explicit procedures around triage based on fire danger indices, fire managers have these indices available and may implicitly use them for triage purposes as fires occur.

In Australia, fire danger indices provide guidance around suppression difficulty of a fire in different fire weather conditions, but do not explicitly indicate fires unlikely to be controlled in initial attack. The McArthur Fire Danger Index for grass (GFDI) and forest (FFDI) was developed to indicate the suppression difficulty of a fire, ranging from low to extreme (and later catastrophic), based on weather and fuel inputs (McArthur 1966, 1967). We refer to these two models combined as FDI, where FFDI is used in forested areas and GFDI in grass. The latest indices developed for the Australian Fire Danger Rating System (AFDRS) use separate models for fire behaviour characteristics and suppression success at initial attack (Hollis, Matthews, Fox-Hughes, *et al*. 2024a), but currently, only the Fire Behaviour Index (FBI) is available for operational use. The AFDRS guides provide descriptions of fire suppression and containment at different FBI values, however these are aligned with fire behaviour with McArthur’s FDI categories (Hollis, Matthews, Anderson, *et al*. 2024b). Therefore, FDI are the most available operational indicator of suppression success likelihood, even though these indices do not directly give likelihood of initial attack success.

There are many published approaches which do address the likelihood of successful initial attack. One option is coupled fire spread and suppression models, beginning with Fried and Fried (1996) and continued in examples such as Finney *et al*. (2008), Tolhurst *et al*. (2008) and Belval *et al*. (2019). These types of models predict the rate of fire area growth, moderated by the rate of suppression applied progressively along the fire edge as determined by a construction rate. However, substantial computing power and data is required for these models to accurately reflect fire and suppression conditions (Granda *et al*. 2023), and the lack of conclusive evidence on suppression rates also limits their ability (Duff and Tolhurst 2015). Therefore, coupled fire spread and suppression models are difficult to implement into fire operations for a rapid triage process.

Empirical models, developed from data on previous fire outcomes, offer a more viable alternative for a rapid assessment. A common approach is to model initial attack success probability using logistic regression with variables such as fuel, weather, landscape terrain, suppression response (e.g. Arienti *et al*. 2006, Plucinski 2012, Plucinski 2013). More recent studies have also examined alternative statistical modelling methods, such as random forest models (Collins *et al*. 2018; Marshall *et al*. 2022). The focus of these studies has been to examine the factors influencing initial attack success or failure, rather than to build a predictive model for operational use. As such, these models include variables such as number of firefighters or delay in response, which are not known prior to the fire starting. There have been some empirical initial attack models developed specifically for an operational context, such as models for Australian forest (Plucinski 2012) and grass fires (Plucinski 2013). Empirical models have been improved over time with the inclusion of more data from expanded, state-wide fire datasets (Wheatley *et al*. 2022; Plucinski *et al*. 2023).

Despite the continued development of these empirical initial attack models, there are challenges to deploying these models into an operational triage process. Some models require data inputs that are unavailable prior to the fire occurring, since they were not developed for this purpose (Collins *et al*. 2018; Marshall *et al*. 2022). Model performance also limits their utility: both the Marshall *et al*. (2022) and Plucinski *et al*. (2023) models predicted incorrect outcomes for 16 37% of fires escaping initial attack in their test datasets, which would mean missing many fires which should have received an elevated response. These studies also do not provide a comparison of new model performance against existing models and tools used operationally, such as fire danger indices. Finally, we also observe very few case study demonstrations of these types of models. These models are critical on days when there are multiple fires competing for resource. Case studies demonstrating the models’ performance and limitations on these days would provide greater certainty around their applicability for an operational context.

This study aims to address these challenges by developing empirical initial attack models with improved performance and providing a more thorough evaluation relative to their intended application. Building on a recent initial attack model (Plucinski *et al*. 2023), we use similar model variables but base the models on a newly curated dataset (Butler *et al*. 2025a) which addresses some of the issues raised in their study around self-contained fires and matching fires in different jurisdictions. We choose variables from operationally available data sources and explore a wider range of modelling techniques with consideration to the interpretability of the models.

For model evaluation, we depart from previous studies focussing on typical performance across all fires by assessing performance on the intended application of the models. We compare our models against base models using the existing metric of FDI and evaluate its performance on days in Victoria’s fire history where ignition triage was required. We expect our models will perform better at discerning where fires are likely to be uncontrolled in initial attack compared to FDI, since the models predict the initial attack outcome directly. By assessing the models performance on case studies of days in Victoria’s fire history where fire triage was required, we expect to gain insights on the limitations and advantages of the models in different circumstances. Overall, we expect that our approach to development and validation for the specific application will yield models suitable for operational use for triaging fires.

## Methods

Our approach to development and validation had five steps, detailed in this methods section. First, we selected data by obtaining a dataset of spreading wildfire incidents for Victoria, Australia, and classified the incidents by initial attack outcome (controlled or uncontrolled). We then selected candidate modelling variables based on availability of information at fire report time and previous studies into factors affecting initial attack outcome (e.g. Collins *et al*. 2018; Marshall *et al*. 2022; Plucinski *et al*. 2023). Our model development involved trialling model types and choosing the best model type based on performance and application requirements. We also undertook a variable selection process. Finally, we evaluated our models by assessing their performance on a test set, comparing their performance against models using only FDI and reviewing their performance on four critical days in Victoria’s fire history.

### Data selection

In this study, we used the Victorian Wildfires Research Dataset (VWFRD) version 2.5.8 (Butler *et al*. 2025a). VWFRD is a tabular dataset of reported unplanned wildfires in regional Victoria, Australia from 1 April 2008 to 30 March 2024, a total of 132,537 records. It contains estimated ignition points, final fire size, reported time and time the fire was declared ‘under control’. The dataset includes data from fires on both private and public land, and resolves duplicated reports across different jurisdictions. An issue with using a dataset of all reported wildfire incidents is that the dataset includes many non-spreading fires that are self-contained by weather and fuel discontinuities, such as unattended campfires and small fires in garden beds or traffic medians (McCarthy *et al*. 2023). This is a problem for initial attack modelling because these non-spreading fires may have similar data values to other fires but will invariably be controlled in initial attack. To resolve this, we filter our data to only the ‘spreading’ fires in the VWFRD with valid locations and times (*n* = 12,357). This is a departure from previous modelling (e.g. Plucinski *et al*. 2023), which uses all fire reports, regardless of whether the fire was spreading, due to not having this information available.

### Initial attack outcome

Our study estimates the probability of unsuccessful initial attack, which can be measured by area (spatially) and by time (temporally): the fire area exceeds a certain size, or the fire is not controlled within a certain timeframe (Korkola *et al*. 2024). Since fires in grass and forest have notable differences in nature of spread, intensity, accessibility and consequence, we chose different thresholds for success based on the main fuel type of the fire. We defined fires as uncontrolled in initial attack in four ways:

- Grass: containment time greater than 2 hrs,
- Grass: final fire area larger than 100 ha,
- Forest: containment time greater than 4 hrs,
- Forest: final fire area larger than 5 ha,

These thresholds represent how grass fires generally spread faster and so burn more area initially but take less time to control. The forest area thresholds align with fire agency operational targets and systems (Country Fire Authority 2020; Department of Energy, Environment and Climate Action 2025).

These choices of outcomes mean our data is split into grass and forest fires. The main fuel type of the fire is determined by the fuel type with the highest density in the 3 km surrounding the fire’s reported location, see Table 1. Shrub and no fuel fires (8% of the total data) were ignored for this study. Therefore, we fit four models: two datasets (grass and forest fires), each with two outcomes (spatial and temporal). See Tables 2 and 3 for details of these datasets. Most of the fires in our datasets were controlled fires, meaning the dataset is unbalanced, see Table 3.

**Table 1.** Modelling variables with description.

| Variable | Units | Description |
| --- | --- | --- |
| **Weather** |  |  |
| **Temperature** | °C | Ambient air temperature, either the most recent forecast before the fire report time from the Australian Digital Forecast Data (ADFD) (Bureau of Meteorology 2024a) or the VicClim grid value closest to the reported fire time and location. |
| **Relative humidity** | % | Relative humidity as a percentage, either the most recent forecast before the fire report time from the ADFD or the VicClim grid value closest to the reported fire time and location. |
| **Wind speed** | km/h | Wind speed on the hour, either the most recent forecast before the fire report time from the Bureau of Meteorology (BoM) or the VicClim grid value closest to the reported fire time and location. |
| **Drought factor** | - | Drought factor, either the most recent forecast before the fire report time from the ADFD or the VicClim grid value closest to the reported fire time and location. |
| **KBDI** | - | Keetch Byram Drought Index, either the most recent forecast before the fire report time from the ADFD or the VicClim grid value closest to the reported fire time and location. |
| **Soil moisture content** | % | Root zone soil moisture (percentage of available water content in the top 1 m of soil) from the closest time and location in the Australian Water Outlook (Bureau of Meteorology 2024b) |
| **Curing** | % | Percentage of dead fuel material within grassland, based on the latest modelled grid value at the the reported fire time and location (Country Fire Authority 2024b). |
| **VicClim indicator** | - | Binary indicator which is 0 when the weather data is forecast ADFD data (2019-2024) and 1 when it is from reanalysis data VicClim (2008-2019). |
| **Fuel** |  |  |
| **Grass density (3km) <sup>A</sup>** | % | Percentage of (30m by 30 m) grid cells that are categorised as grass in the operational fuel layer (Forest Fire Management Victoria 2021) in a grid with the fire start location at the centre and apothem 3 km. |
| **Forest density (3km) <sup>A</sup>** | % | Percentage of (30m by 30 m) grid cells that are categorised as forest by a fuel layer (Forest Fire Management Victoria 2021) in a grid with the fire start location at the centre and apothem 3 km. |
| **Shrub density (3km) <sup>A</sup>** | % | Percentage of (30m by 30 m) grid cells that are categorised as shrub by a fuel layer (Forest Fire Management Victoria 2021) in a grid with the fire start location at the centre and apothem 3 km. |
| **No fuel density (3km) <sup>A</sup>** | % | Percentage of (30m by 30 m) grid cells that are categorised as no fuel by a fuel layer (Forest Fire Management Victoria 2021) in a grid with the fire start location at the centre and apothem 3 km. |
| **Distance to grass-bush interface** | m | Manhattan distance, counting the number of 30 m grid cells to the boundary between the grass and bush (forest or shrub). If the fire start location is in grass, it is the distance to bush and vice versa. If the fire starts in no vegetation, the value is set to zero. Calculated from a vegetation layer (Forest Fire Management Victoria 2021). |
| **Topography** |  |  |
| **Elevation** | m | Distance above sea level, 30 m grid (Vicmap 2024a). |
| **Ruggedness average (3 km)** | - | Average ruggedness in a 3 km apothem square around the fire start location. Ruggedness calculated according to Reily *et al.* 1999. |
| **Access and response** |  |  |
| **Building density (3 km)** | buildings | Number of buildings within a 3 km radius of the fire start location. This variable captures whether the fire started in a town or not. Calculated from Vicmap buildings data (Vicmap 2024b). |
| **Building density (20 km)** | buildings | Number of buildings within a 20 km radius. This variable captures the remoteness of the fire, as more buildings in a 20 km radius means there is likely to be a faster response time. Calculated from Vicmap buildings data (Vicmap 2024b). |
| **Road density (3 km)** | km | Distance of road in kilometres in a square of apothem 3 km around the fire report location. Calculated from Vicmap transport road line data (Vicmap 2024c). |
| **Distance to road** | m | Geodesic distance to the nearest road from the fire report location. Calculated from Vicmap transport road line data (Vicmap 2024c). |

**Table 2.** Summary of dataset split by fuel type.

| **Fuel** | **Count of fires** | **Percentage of total fires (%)** |
| --- | --- | --- |
| **Grass** | 7,580 | 61 |
| **Forest** | 3,812 | 31 |
| **Shrub** | 443 | 4 |

<sup>A</sup> The four vegetation density variables add to 100%, so at most three out of the four are used in any model. For example, for the grass model, grass is considered the baseline and is not included in the modelling variables.

|  |  |  |
| --- | --- | --- |
| **No fuel** | 522 | 4 |
| **Total** | 12,357 | 100 |

**Table 3.** Summary of datasets used for the four regression models and their outcomes.

| **Fuel** | **Outcome type** | **Initial attack outcome** | **Binary indicator** | **Count** | **Percentage** |
| --- | --- | --- | --- | --- | --- |
| **Grass** | Temporal | Controlled - Containment time less than 2 hours | 0 | 6,584 | 87 |
|  |  | Uncontrolled - Containment time greater than 2 hours | 1 | 996 | 13 |
|  |  | Total |  | 7,580 | 100 |
|  | Spatial | Controlled - Final fire area less than 100 ha | 0 | 7,362 | 97 |
|  |  | Uncontrolled - Final fire area greater than 100 ha | 1 | 218 | 3 |
|  |  | Total |  | 7,580 | 100 |
| **Forest** | Temporal | Controlled - Containment time less than 4 hours | 0 | 2,821 | 74 |
|  |  | Uncontrolled - Containment time greater than 4 hours | 1 | 991 | 26 |
|  |  | Total |  | 3,812 | 100 |
|  | Spatial | Controlled - Final fire area less than 5 ha | 0 | 3,237 | 85 |
|  |  | Uncontrolled - Final fire area greater than 5 ha | 1 | 575 | 15 |
|  |  | Total |  | 3,812 | 100 |

### Model input variables

We followed four principles when selecting candidate modelling variables, listed in Table 1:

1. Terrain variables should be robust to errors in the reported start location of the fire.
2. Terrain variables should account for the area the fire spreads into.
3. Weather variables should represent the latest forecasted weather available at the fire report time.
4. Suppression response resources should not be included in the model.

Terrain variables must be robust to errors in the reported location of the fire, because the reported location is not always the fire’s ignition point. For example, private land fires are often reported at the property address point of the fire. Phan and Kilinc (2015) found that the average distance between reported location and point origin in a subset of records in Victoria was 3.7 km. To account for this inaccuracy, we aggregate terrain attributes in an area surrounding the reported fire location. For example, we measure ruggedness by taking the average of the ruggedness of all grid points within a square of apothem 3 km centred on the fire location (see Table 1). Defining model variables in an area around the fire start location also serves the second principle, because it incorporates information about the terrain around the fire.

Our models use the Bureau of Meteorology’s Australian Digital Forecast Database (ADFD) (Bureau of Meteorology 2023) forecast weather variables between 2019-2024 and reanalysis VicClim weather between 2008-2019 (described in Brown *et al*. 2016, updated 2020), since neither dataset covers the full period of our study. The ADFD would be input into the model for use operationally, since it provides the latest forecast for a grid across the state, so in the period where the data overlaps, we have used the ADFD. This approach is in alignment with Kenny *et al*. (2024).

Suppression resources were not included in the models, since we intend the models to help with decisions to escalate resources. Therefore, the models only include information that is known at the time of the fire being reported.

These four principles were implemented in the candidate modelling variables shown in Table 1 to produce the modelling dataset (Butler *et al*. 2025b). The data fall into four groups: weather, fuel, topography, and access and response.

### Modelling

The principle guiding our modelling was that the models should be as simple and explainable as possible so that decisions based on them are transparent and justifiable. The modelling was done in four parts: splitting the dataset into test and training data, choosing a modelling technique, selecting variables, and refitting the models (Fig. 1).

**Fig. 1.**
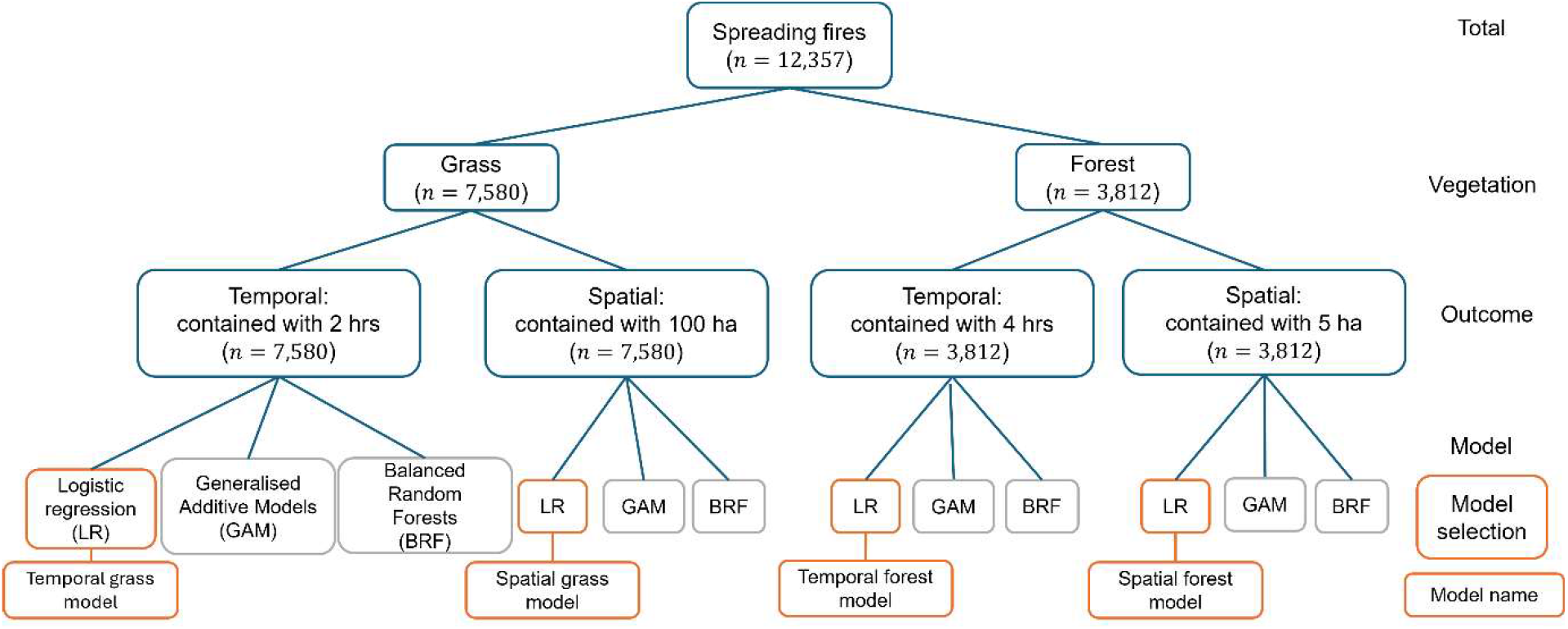
Tree diagram representing the twelve models fit in our analysis. It demonstrates how the full dataset of spreading fires was first split by vegetation type, and then each vegetation type was assigned a temporal and spatial outcome. Finally, each outcome was modelled via logistic regression, generalised additive models and balanced random forests. After model selection (indicated in orange), the four logistic regression models were chosen for detailed analysis and given names. The rest of the models (indicated in grey) are detailed in the supplementary materials.

We split the data into training data (to fit the models) and testing data (to assess their performance on new data) by random sampling, with two considerations:

1. Similar fires often occur on the same day in similar locations, so randomly distributing them across the train and test sets might overestimate the models performance.
2. There were two fire seasons with major fires in our data (2008-09 and 2019-20, (Country Fire Authority 2024a)), and it might over (or under) estimate the model performance if both of those years were included in the training (or test) set by randomisation.

We first randomly allocated one of the fire seasons with major fires to the training set and the other to the test set. We then split the rest of the data by randomly sampling 70% of fire seasons into the training set and 30% into the test set (leading to roughly 80%-20% split across the data), see Supplementary material S1 for further details.

We tested logistic regression, balanced random forests and generalised additive models (GAMs). In Python, we used statsmodels (Seabold and Perktold 2010) to fit logistic regression, and sklearn (Pedregosa *et al*. 2011) and imblearn (Lemaître *et al*. 2017) to fit balanced random forests. The GAMs were fit in R using mgcv (Wood 2011, 2017). All the methods gave similar results, so we chose logistic regression as our preferred technique since it was the simplest and most explainable method (Supplementary material S2 and S4). We used randomised quantile residual plots (Dunn and Smyth 2018) to check the linearity assumption of logistic regression and found it to hold (Supplementary material S3). We applied a log(*x* + 1) transform to variables that appeared skewed in the residual plots, giving our final candidate variables.

Variables were selected for the final models using adaptive Lasso (Zou 2006). We tested for multicollinearity using the variance inflation factor, which was found to be low (less than five) for all the variables in both grass and forest datasets. We then selected variables using adaptive Lasso (Tibshirani 1996; Zou 2006) with the Bayes Information Criterion to choose the penalisation parameter (Wang *et al*. 2007; Zhang *et al*. 2010) via the glmnet package in R (Friedman *et al*. 2010; Tay *et al*. 2023).

### Model evaluation

We evaluated the models performance in three ways: assessing the model performance on the test dataset, comparing the model to a base model using a fire danger index and investigating performance on historical case study days. For the base models, we developed logistic regression models for grass and forest fires using GFDI and FFDI as the only input variable to the model respectively. The FDI values were calculated at the 1.5 km grid cell level, for the fire’s reported location and hour of report. This level of information is available to fire managers operationally. There are no specific FDI thresholds employed by fire managers for triage purposes to benchmark our initial attack models against, so we modelled FDI in a logistic regression model as the outputs are directly comparable to the new models.

For the case study evaluation, we selected four days in Victoria’s fire history where resource triage was required, shown and described in Table 4. For the evaluation, we re-trained the grass and forest temporal models on the full dataset. Therefore, the test data is included in the training data for the case study evaluation. We reviewed the combined performance of the grass and forest temporal models over each of the days, looking at the models ability to distinguish between spreading fires at times where there are multiple concurrent fires, potentially requiring triage for an escalated response. We investigated specific incidents in detail where the models outputs were unexpected compared to the actual fire outcome, consulting situation reports and radio logs, to identify any limitations of the modelling.

**Table 4.** Case study days selected for testing model performance.

| Day | Description | Grass |  |  | Forest |  |  |
| --- | --- | --- | --- | --- | --- | --- | --- |
|  |  | Initial attack outcome (temporal) |  | Total | Initial attack outcome (temporal) |  | Total |
|  |  | Controlled | Uncontrolled |  | Controlled | Uncontrolled |  |
| **7 February 2009**<br>**Black Saturday** | The most fatal wildfire event in Victoria's recent fire history, which led to 173 deaths and 400,000 ha burned from 13 significant fires (Country Fire Authority 2023). | 3 | 9 | 12 | 3 | 12 | 15 |
| **17 March 2018**<br>**St Patricks Day**<br>**Fires** | Strong gusts in south-west Victoria ahead of a cold front caused electricity infrastructure failures to ignite a series of grass fires late in the evening. | 11 | 2 | 13 | 3 | 1 | 4 |
| **21 November 2019**<br>**Black Summer**<br>**Ignitions** | Major day of ignitions for Australia's 'Black Summer' season 2019-20, where forest campaign fires eventually burnt over 7 million hectares in South-Eastern Australia (Australian Institute for Disaster Resilience 2020). Three of the most significant fires for this season were ignited by a series of lightning strikes in inaccessible and rugged terrain on 21 November 2019, which grew under strong winds and worsening overnight conditions. | 8 | 4 | 12 | 8 | 20 | 28 |
| **17 February 2023** | A more recent example of a typical fire season day in 2023 (17 February), where two grass fires were uncontrolled in initial attack and triage of aircraft between fires was required. | 11 | 2 | 13 | 3 | 1 | 4 |

## Results

### Model performance

To assess the performance of the four models (see Tables 5 to 8), we plotted the probability distribution of the model predictions on the test data, split by the ground truth: whether the fires were controlled or uncontrolled in initial attack. We refer to these as *separation plots*, because a well-performing model should show the distribution of controlled fires (blue) to the left and of uncontrolled fires (orange) to the right.

**Table 5.** Coefficients and model statistics for the temporal initial attack model for grass fires. The model estimates the probability of being uncontrolled (containment time will be greater than 2 h).

|  | **Coefficient** | **std error** | **p-value** |
| --- | --- | --- | --- |
| **Intercept** | -3.7005 | 0.635 | 0.000 |
| **Temperature** | 0.0129 | 0.009 | 0.172 |
| **Relative humidity** | -0.0062 | 0.003 | 0.054 |
| **log Wind speed** | 0.3689 | 0.086 | 0.000 |
| **KBDI** | 0.0035 | 0.001 | 0.003 |
| **log Soil moisture content** | -0.0035 | 0.044 | 0.937 |
| **Curing** | 0.0096 | 0.003 | 0.000 |
| **VicClim indicator** | 0.3493 | 0.115 | 0.002 |
| **Forest density (3 km)** | 0.0109 | 0.004 | 0.003 |
| **log Shrub density (3 km)** | 0.1768 | 0.039 | 0.000 |
| **log Ruggedness average (3km)** | 0.3677 | 0.059 | 0.000 |
| **log Building density (3 km)** | -0.1775 | 0.048 | 0.000 |
| **log Road density (3 km)** | -0.4889 | 0.111 | 0.000 |
| **log Distance to road** | 0.2986 | 0.023 | 0.000 |

**Table 6.** Coefficients and model statistics for the spatial initial attack model for grass fires The model estimates the probability of being uncontrolled (final fire area will be greater than 100 ha).

|  | **Coefficient** | **std error** | **p-value** |
| --- | --- | --- | --- |
| **Intercept** | -8.4225 | 1.139 | 0.000 |
| **Temperature** | 0.0702 | 0.011 | 0.000 |
| **log Wind speed** | 0.9663 | 0.177 | 0.000 |
| **Drought factor** | 0.0902 | 0.060 | 0.134 |
| **Curing** | 0.0303 | 0.007 | 0.000 |
| **log No veg density (3km)** | -0.5678 | 0.136 | 0.000 |
| **log Distance to grass-bush interface** | 0.2801 | 0.092 | 0.002 |
| **log Ruggedness average (3km)** | 0.3770 | 0.106 | 0.000 |
| **log Building density (20 km)** | 0.0414 | 0.071 | 0.559 |
| **log Road density (3 km)** | -0.9260 | 0.214 | 0.000 |
| **log Distance to road** | 0.1639 | 0.043 | 0.000 |

**Table 7.**
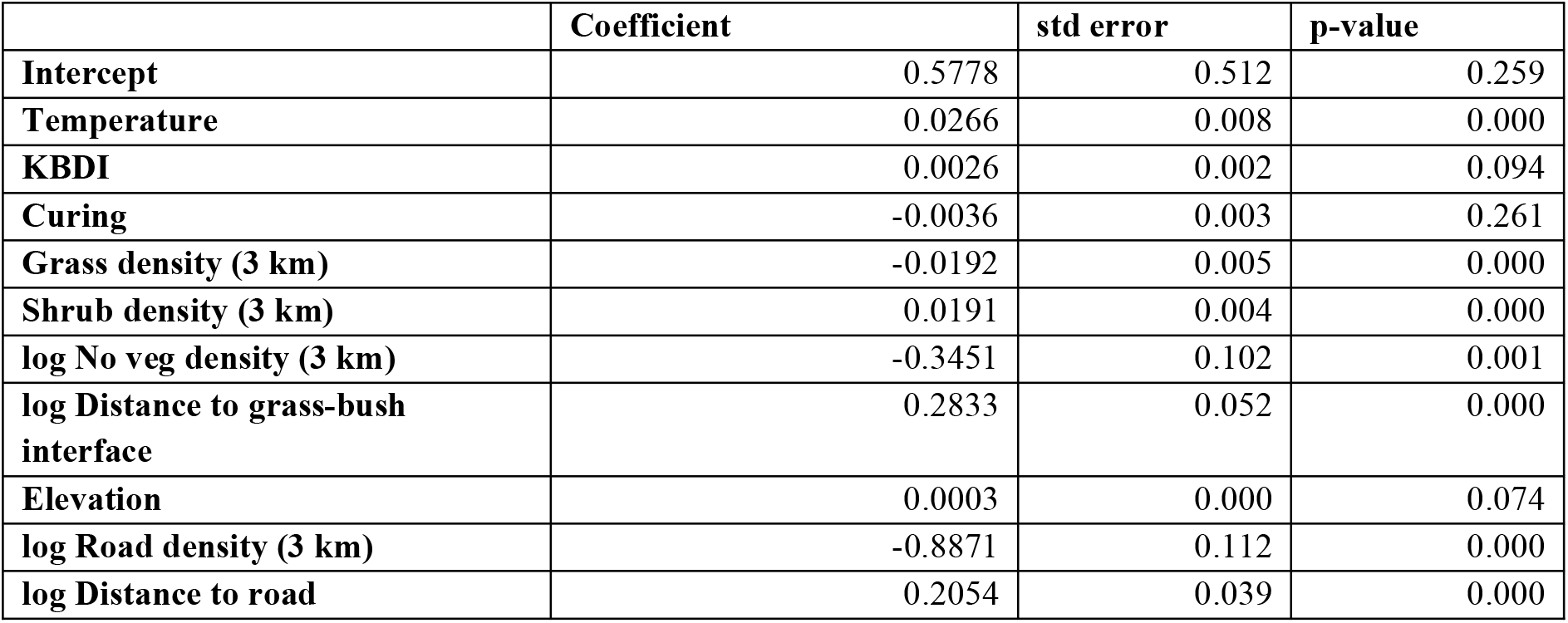
Coefficients and model statistics for the temporal initial attack model for forest fires The model estimates the probability of being uncontrolled (containment time will be greater than 4 h).

|  | **Coefficient** | **std error** | **p-value** |
| --- | --- | --- | --- |
| **Intercept** | 0.5778 | 0.512 | 0.259 |
| **Temperature** | 0.0266 | 0.008 | 0.000 |
| **KBDI** | 0.0026 | 0.002 | 0.094 |
| **Curing** | -0.0036 | 0.003 | 0.261 |
| **Grass density (3 km)** | -0.0192 | 0.005 | 0.000 |
| **Shrub density (3 km)** | 0.0191 | 0.004 | 0.000 |
| **log No veg density (3 km)** | -0.3451 | 0.102 | 0.001 |
| **log Distance to grass-bush interface** | 0.2833 | 0.052 | 0.000 |
| **Elevation** | 0.0003 | 0.000 | 0.074 |
| **log Road density (3 km)** | -0.8871 | 0.112 | 0.000 |
| **log Distance to road** | 0.2054 | 0.039 | 0.000 |

**Table 8.** Coefficients and model statistics for the spatial initial attack model for forest fires. The model estimates the probability of being uncontrolled (final fire area will be greater than 5 ha).

|  | Coefficient | std error | p-value |
| --- | --- | --- | --- |
| **Intercept** | 0.2425 | 0.611 | 0.691 |
| **Relative humidity** | -5.0151 | 0.003 | 0.000 |
| **Drought factor** | 0.0934 | 0.032 | 0.003 |
| **log Wind speed** | 0.2355 | 0.110 | 0.033 |
| **Elevation** | 0.0002 | 0.000 | 0.235 |
| **log Building density (20 km)** | -0.2635 | 0.040 | 0.000 |
| **log Road density (3 km)** | -0.5067 | 0.104 | 0.000 |
| **log Distance to road** | 0.0877 | 0.038 | 0.021 |
| **Grass density (3 km)** | -0.0107 | 0.004 | 0.014 |

Across all four models, separation was poor, suggesting limited ability to distinguish controlled from uncontrolled fires (Fig. 2a to Fig. 5a). The distributions of both the controlled and uncontrolled fires are right-skewed, meaning the model predicts most fires to have a low probability of being uncontrolled. This is understandable, since most of the fires in the dataset are controlled. However, it does indicate that our input variables are not sufficient to distinguish well between controlled and uncontrolled fires. The area under the curve and average precision metrics further support these conclusions (Table 9). Further model diagnostics that support these conclusions and aid interpretation of the separation plots are given in Supplementary material S2.

**Table 9.**
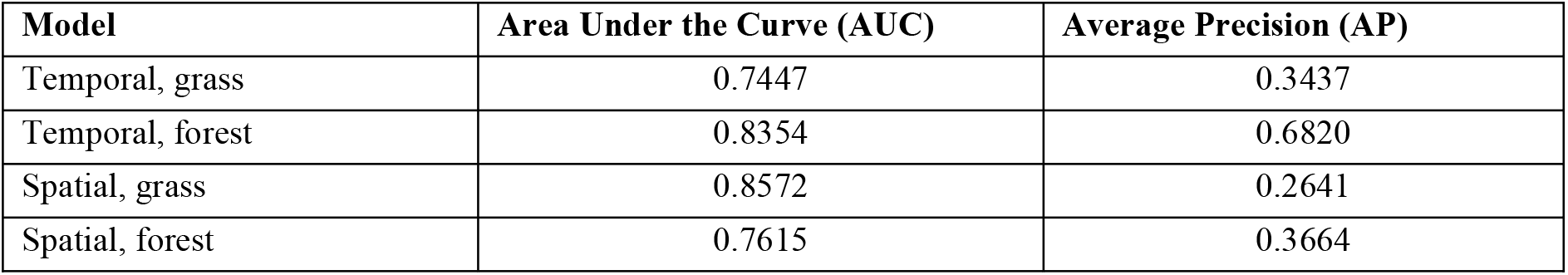
Model performance metrics for the four initial attack models. The AUC values seem fairly high, but are not an accurate representation of performance due to the unbalanced response (many more controlled fires than controlled). The AP measure gives a better indication, which is consistent with the conclusion from the separation plots that the temporal, forest model performs best.

**Fig. 2.**
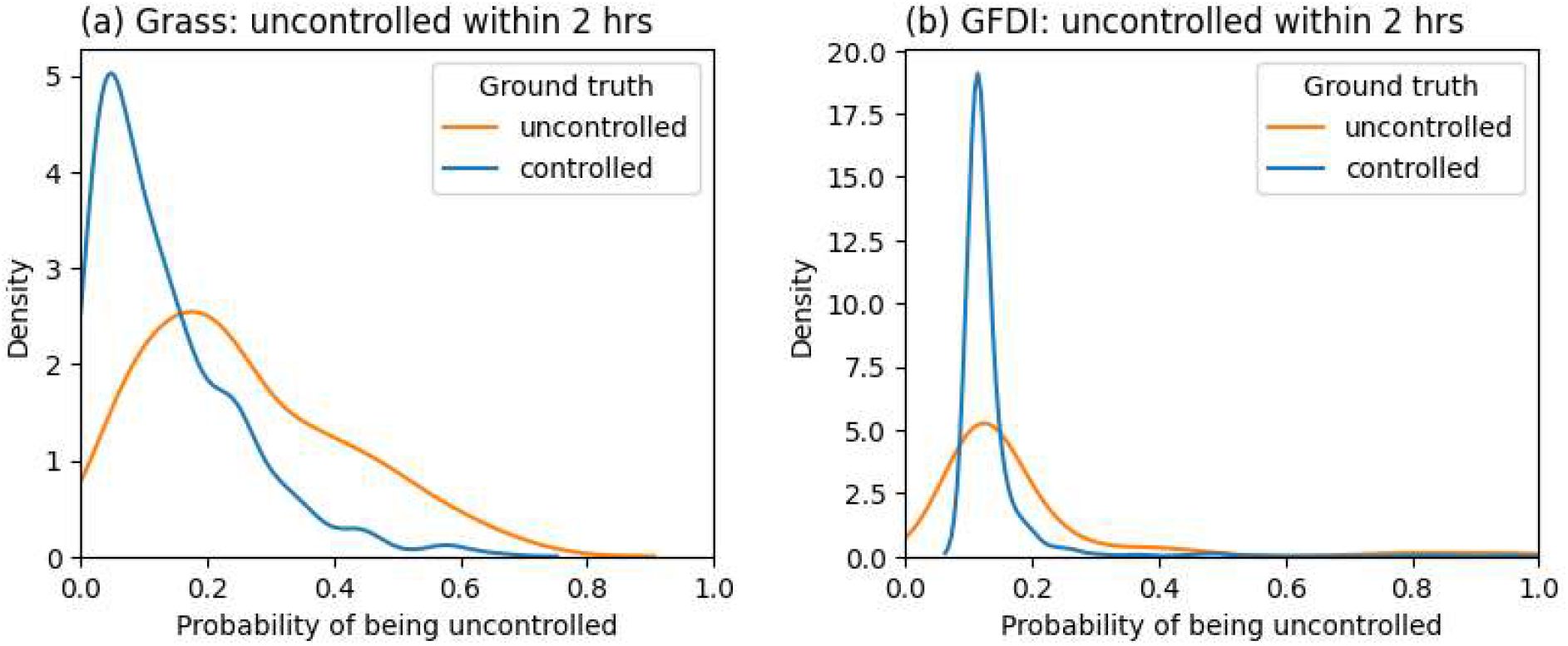
Separation plot for the model (a) compared to a base model only using GFDI (b) for grass fires, estimating the probability of being uncontrolled (containment time will be greater than 2 h). The model has poor separation but is distinguishing between controlled (blue) and uncontrolled (orange) fires better than the base model.

The best performance is by the temporal forest model. Despite poor separation, the separation plot in Fig. 3a indicates that when the probability of being uncontrolled is higher than 0.5, most of the fires truly are uncontrolled.

**Fig. 3.**
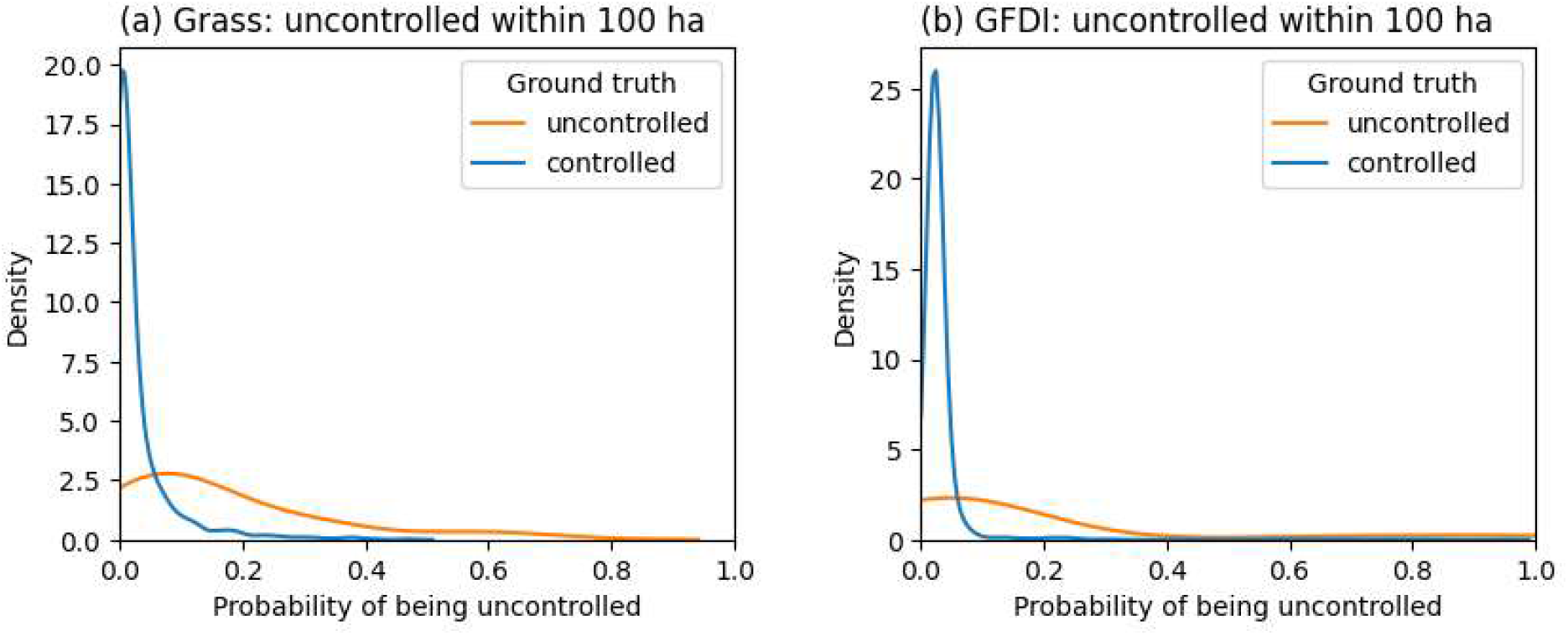
Separation plot for the model (a) compared to a base model only using GFDI (b) for grass fires, estimating the probability of being uncontrolled (final fire area will be greater than 100 ha). The model has poor separation but is distinguishing between controlled (blue) and uncontrolled (orange) fires slightly better than the base model.

**Fig. 4.**
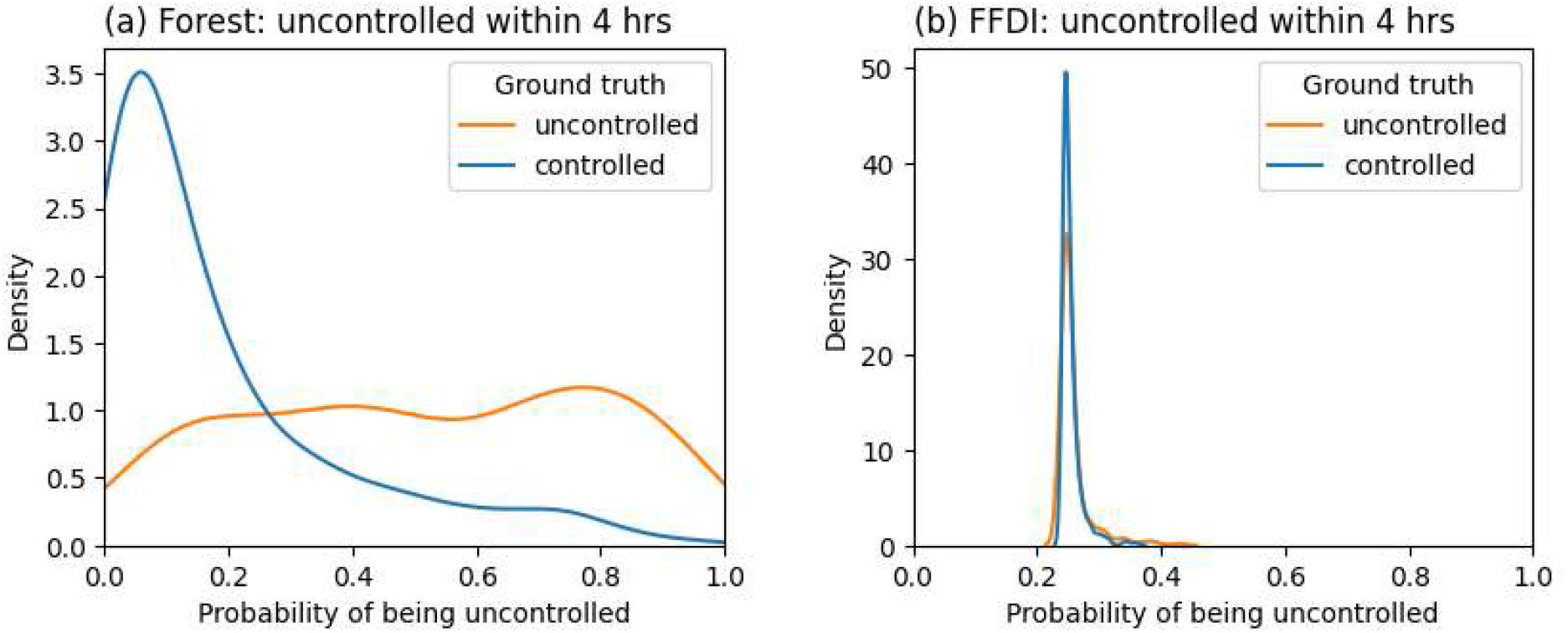
Separation plot for the model (a) compared to the base model only using FFDI (b) for forest fires, estimating the probability of being uncontrolled (containment time will be greater than 4 h). The model has some separation but is distinguishing between controlled (blue) and uncontrolled (orange) fires significantly better than the base model.

### Comparison with base models

We also compared our models to base models using only the grass fire danger index (GFDI) or forest fire danger index (FFDI) (Fig. 2b to Fig. 5b). The diagrams show very poor separation between controlled and uncontrolled outcomes for both the GFDI and FFDI models. This demonstrate that our initial attack models distinguish better than FDI for both grass and forest fires.

**Fig. 5.**
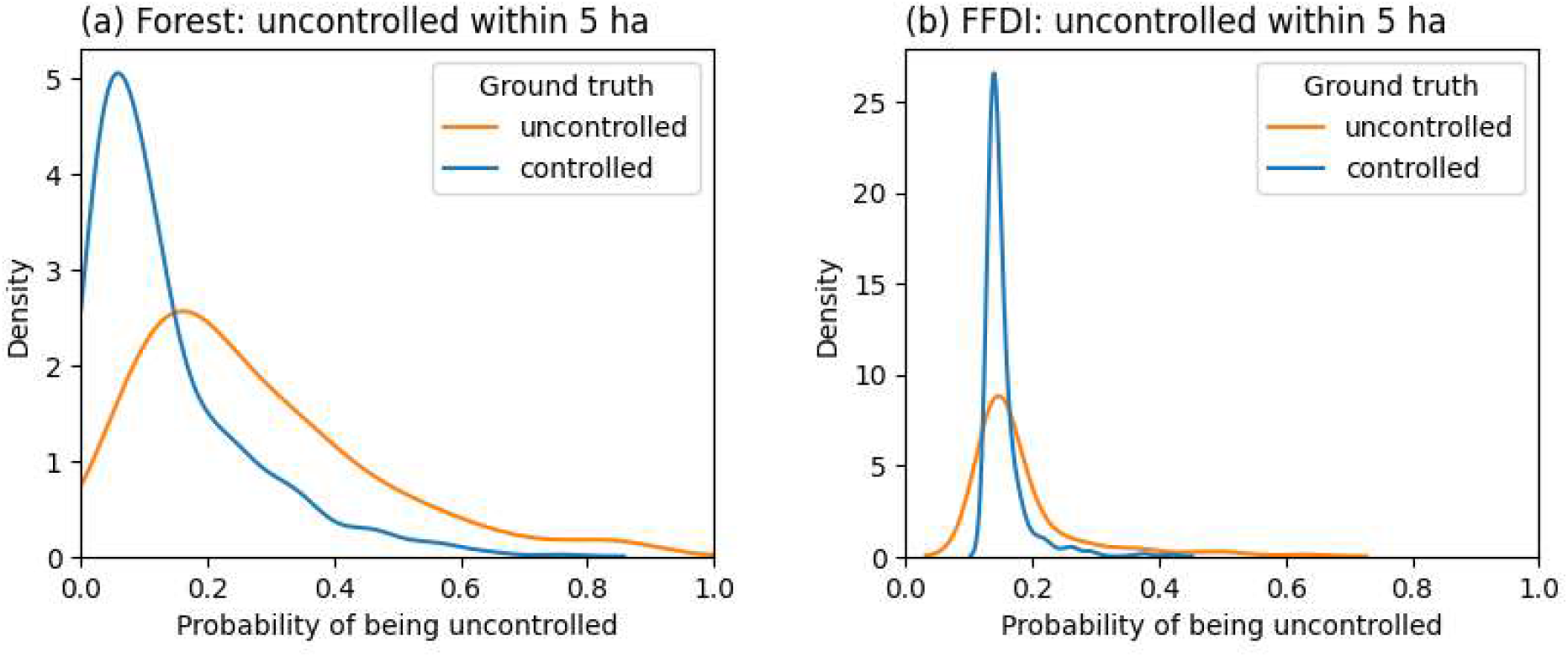
Separation plot for the model (a) compared to the base model only using FFDI (b) for forest fires, estimating the probability of being uncontrolled (final fire area will be greater than 5 ha). The model has poor separation but is distinguishing between controlled (blue) and uncontrolled (orange) fires better than the base model.

### Case studies

The probability of the spreading fires being uncontrolled, as predicted by our temporal outcome models, is shown over the course of the day for the four case study days (Fig. 6). Good model performance would be indicated on these plots by controlled fires (shown in blue) being clustered at the bottom of the plot and uncontrolled (shown in orange) clustered at the top of the plot. In these plots, we observe mixed results: most fires are allocated probabilities below 0.5, with mixed ground truth outcomes (uncontrolled and controlled). Most of the fires with higher probabilities were uncontrolled. This is a similar pattern to the separation plots.

**Fig. 6.**
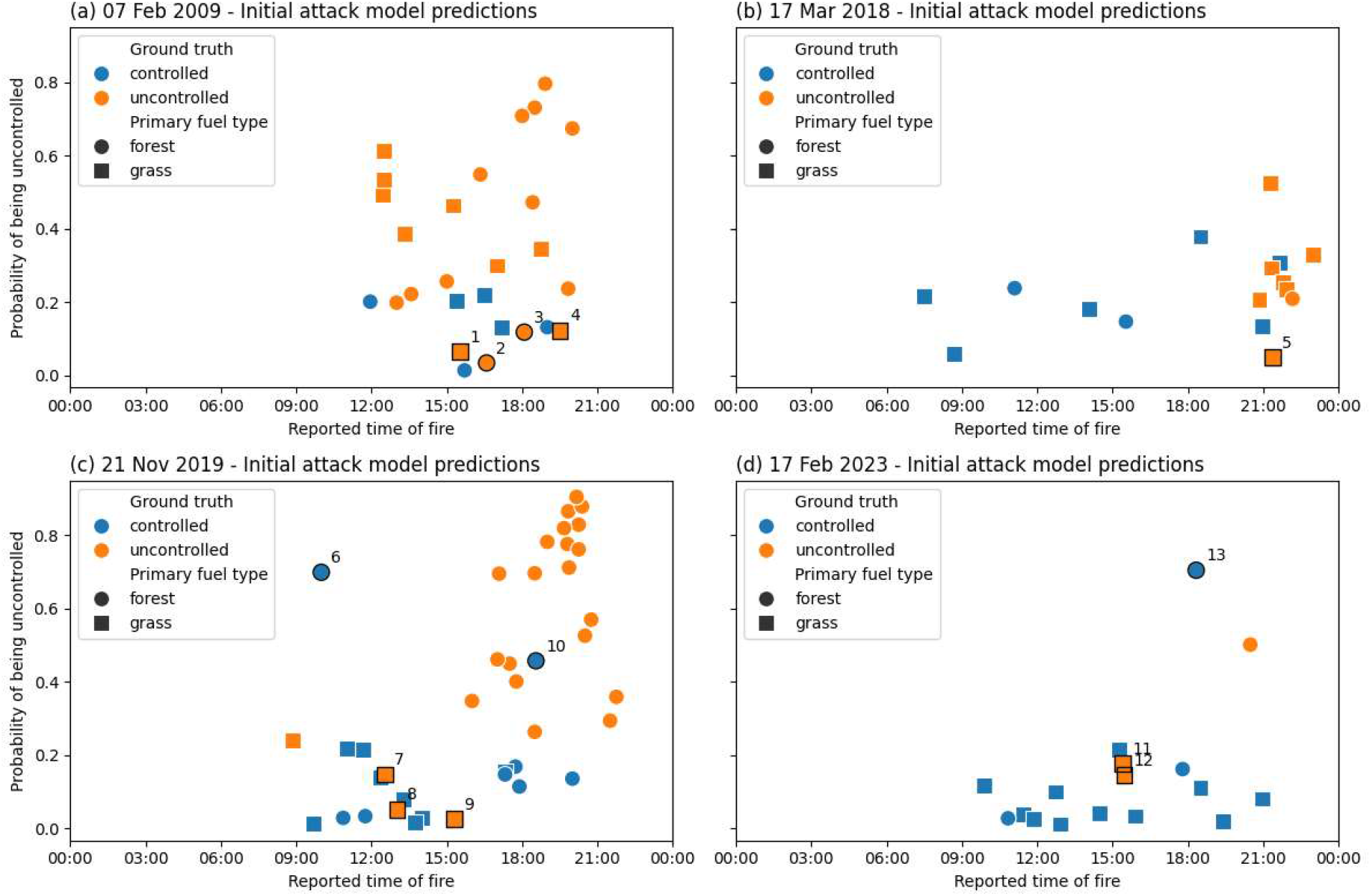
Model predictions for reported grass and forest fires on fires, c) Black Summer lightning ignitions and d) 17 February 2023. Probabilities are reported for both forest (circles) and grass (squares) fires. Actual initial attack outcome is indicated by colour; blue fires were controlled in initial attack and orange fires were uncontrolled. Fires with black outlines and numbered were investigated further.

There are two instances in the case studies where there are many concurrent ignitions, where a model may be useful to assist with triage. The first, after 20:30 on 17 March 2018 (Fig. 6b) shows ineffective separation of uncontrolled and controlled fires with the grass model, returning mostly low probabilities for all fires. The second instance, after 16:00 on 21 November 2019 (Fig. 6c), shows effective separation of fires with the forest model. All uncontrolled incidents have probabilities above 0.2, and there is only one controlled incident with a probability higher than this value. This means with a probability threshold of 0.2, the model would correctly distinguish all but one of the fires as controlled or uncontrolled.

In Fig. 6, we visually identified fires where the model probabilities are unexpected compared to the ground-truth outcome and conducted a more thorough review to identify any notable factors influencing initial attack. We split these into missed detection errors, where the model gives a relatively low probability but initial attack was unsuccessful (fires 1 - 5, 7 - 9, 11 and 12 in Fig. 6, and false alarm errors, where the model gives a relatively high probability of being uncontrolled but initial attack was successful (fires 6, 10 and 13 in Fig. 6). A summary of the review is shown in Table 10, with more details given for each fire in Supplementary material S5.

**Table 10.**
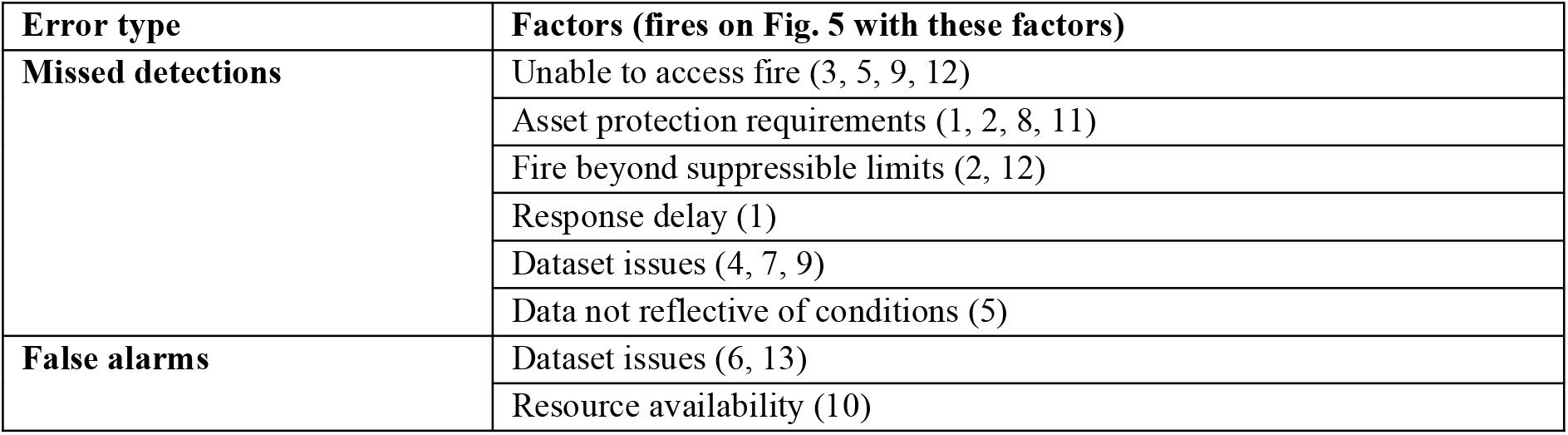
Factors leading to unexpected model results compared to initial attack outcomes based on the review.

For the missed detection errors, we found that the initial attack outcome was influenced by factors not modelled that prevented a standard initial attack response. The requirement to undertake asset protection, as houses or sheds were immediately under threat by fire, prevented firefighting resources from undertaking suppression activities in four of the missed detection instances. In four cases, firefighting resources were unable to safely access the fireground, due to powerlines and hazardous trees. On one of the instances on Black Saturday, resource constraints meant aircraft were unavailable where they normally would have been dispatched. Finally, we found two instances where fire activity was beyond the limits of direct suppression, which prevented standard initial attack actions from occurring.

Dataset issues were found in several of the errors, both false alarms and missed detections. One error stemmed from mislabelling incidents as spreading: Fire 6 was a logging coupe fire that was fully contained at the time of ignition and should not have been included in the model. Misclassification of controlled or uncontrolled in initial attack occurred in two cases. For example, fire 13 was initially recorded as controlled during the initial attack, but situation reports indicated it continued to grow for 24 hours, meaning this was not truly a false alarm. A similar issue was observed in fire 7. Fire 9 should have been classified as a forest fire, where it would have been considered controlled, but it was included in the grass model due to the high presence of surrounding grass. The number and variety of these dataset errors highlight that further work is still required to improve fire datasets for empirical modelling.

## Discussion

We developed initial attack models with the expectation that they would be suitable for use to triage fires for an escalated response. The models were able to better distinguish between initial attack outcomes compared to base models using only GFDI or FFDI, an operational metric available in Victoria indicating suppression difficulty. This is expected due to the addition of new variables related to response and access, fuel and topography, which have been shown to affect initial attack outcome (Collins *et al*. 2018; Rodrigues *et al*. 2019; Marshall *et al*. 2022; Plucinski *et al*. 2023) but are not incorporated into FDI. Our models would therefore be a better candidate than FDI for an operational tool to triage fires.

However, the evaluation suggests that these models are not appropriate for triaging fires on days where resources are limited, which was the intended use case. The models failed to distinguish fires where difficulties such as access, asset protection requirements or communication problems prevented a standard initial attack suppression response, particularly for grass fires. These constraints, or ‘bottlenecks’ preventing or delaying suppression, are highly localised in time and space, making them difficult to be represented by variables which are robust to errors in reported fire location. Given the prevalence of bottlenecks affecting initial attack outcome in the fires reviewed, it is logical to assume they contribute to a significant proportion of uncontrolled fires in the wider dataset. Since the models cannot predict where bottlenecks will occur, their operational usefulness is limited. Additionally, extreme fire behaviour can make direct initial attack impossible to enact safely, as seen on Black Saturday (Teague *et al*. 2010) (see Fig. 6a). In these cases, almost all fires are likely to be uncontrolled in initial attack, so tools that assess the potential consequences and ability to prevent impacts and loss may be more appropriate than initial attack models for triage applications.

We also did not observe an improvement in the model performance using spreading fires, compared to similar models using all fires. Visual comparison of the separation plots shows similar results between our models and (Plucinski *et al*. 2023). Despite the lack of improvement in model performance, we suggest accounting for non-spreading fires in fire reporting datasets is essential for modelling, as they represent a different entity compared to a wildfire with sufficient fuel to develop to steady state. The dataset we used in our model, VWFRD, uses a rule-based procedure to identify spreading fires and is subject to errors, as found in our case studies. Additional work should look at improving this process so that initial attack modelling can focus on fires that require initial attack to be suppressed. We also did not consider how non-spreading fires would fit within an operational triage process: we assumed an application of triaging spreading fires only. A two-step hurdle model that firstly identifies the probability that a reported fire is a spreading fire and then applies an initial attack model could be a more appropriate process.

We observed good model performance on one case study, where the forest temporal model effectively separates initial attack outcomes for remote, lightning-ignited forest fires. In the case study of the 21 November 2019, the model could have assisted accurately triaging fires for an escalated response. This model also showed the best performance overall on the separation plots, potentially from the influence of other, similar lightning ignition events. These lightning ignition events have historically caused significant losses and area burned in Victoria compared to other ignition types (Read *et al*. 2018), and so using the model for triaging fires for this specific circumstance alone could provide some value. Further validation would be required to test the model’s performance on other similar events and before trialling in an operational setting.

Data availability is a limiting factor in improving initial attack models to make them appropriate for triage. We tested more flexible model types, GAMs and random forest, and showed that there was no significant improvement in model skill with new model types, and therefore improvements are likely to come from the inclusion of new variables. This was supported by case studies identifying that the model was missing the influence of initial attack bottlenecks. Variables that described these bottlenecks would violate our modelling principles and, therefore, make the model impractical for operational use. For example, local access, asset locations and detailed fuel attributes required to improve the grass fire model would not be robust to errors in the reported fire location. Response delay is not available to agencies in a systematic way prior to the fire starting. Incorporating resource location and availability metrics, such as fire load (e.g. Wheatley *et al*. 2022) and depot or brigade locations could improve the model and would fit within our modelling principles but will not fully address the missing factors.

Our approach to combine the use of separation plots and case studies resulted in useful findings on the applicability of the models for triaging fires for resource allocation. Previous studies which build similar initial attack models either suggest a use case for triage decision support (Plucinski 2013; Wheatley *et al*. 2023) or suggest general statistical findings can be used by fire managers for allocation of resources (Collins *et al*. 2018; Rodrigues *et al*. 2019). However, these studies do not demonstrate the model applied to the proposed use case. We have shown that case study validation demonstrates a more nuanced view of where models may be useful, demonstrating limitations on its applicability. Our case study set was limited to only four days of testing, and future work could consider a more thorough evaluation on more days to check these findings hold across other examples. These conclusions also cannot necessarily be generalised to other applications, so we encourage a similar evaluation process for initial attack models for use in policy, risk or preparedness.

## Conclusions

We developed and evaluated empirical initial attack models for the specific purpose of triaging fires for an escalated response. We showed that the models perform better than available metrics, but have limited broad skill and as they do not account for cases where bottlenecks prevent a standard initial attack response from occurring. They also may not be appropriate for days with extreme fire behaviour or where resources are constrained. Therefore, the study showed the models are not suitable for operational use for triaging all wildfire ignitions in Victoria. The temporal forest model had the best performance in determining whether a forest fire will be uncontrolled in initial attack (4 h), particularly for lightning ignition events. Future work could include further improvements to that model and trialling its use in an operational setting alongside current metrics.

This work demonstrates the value in developing and validating an initial attack model with a specific use case in mind. The application-specific approach guided decisions on model variables, such as choosing those available operationally prior to the fire starting. We considered interpretability of the models and chose logistic regression ahead of more complex models since they gave no significant gain in performance. Finally, we used case studies to identify limitations of the models performance. We found this approach beneficial compared to a more general study of initial attack success and recommend it is adopted for the development of empirical models for other, related applications.

## Acknowledgements

The authors would like to acknowledge Tom Duff, Tim Gazzard and Matt Plucinski for insightful comments on the manuscript.

## Data availability statement

Data and code used to produce this study is available at https://github.com/vic-ffm/vic-ia. For a more detailed version of the dataset, including fire location and report time, please contact the authors.

## Supplementary material

### Supplementary material S1. Train and test split

As described in the Methods, the data was split into a training set used to fit the model parameters and a test set to assess the model performance. The fire seasons were split such that 76% of the fire seasons are in the training set and 24% in the test set, with each of the train and test set including one season with major fires, see Table S11 for the full details. Since each fire season has a different number of fires, the percentage split in terms of rows of data differs slightly and is given in Table S2.

### Supplementary material S2. Diagnostic plots for logistic regression models

We present some standard diagnostic plots for our logistic regression models. These plots are on the test data set. In each of Fig. S1 S4, the left plot is the same separation plot shown in the main results, the central plot is the Receiver Operator Curve (ROC), and the right plot is the Precision-Recall Curve. Since our data is unbalanced, the Precision-Recall Curve is a better measure of model performance than the ROC, since it measures the model’s ability to correctly identify uncontrolled fires, which are the more important case to correctly identify. The plots show the Area Under the Curve (AUC), which is the area under the ROC. We show this value, since it is a standard metric for model performance, but note that it is not an appropriate metric for unbalanced data. The Average Precision (AP) is considered a better metric for unbalanced data, since it measures how well the model identifies the positive case, which for our model represents uncontrolled fires. All of these metrics give a similar message to the separation plot: the models do not perform well at identifying uncontrolled fires, and the model with the best performance is the temporal forest model.

### Supplementary material S3. Residual plots for logistic regression models

To assess whether the linearity assumption holds well enough for our data, we used randomised quantile residual plots (Dunn and Smyth 2018). Standard residual plots for logistic regression are uninformative, since the binary outcomes do not produce a plot that is visually easy to diagnose. Randomised quantile residual plots overcome this problem by creating a plot which is visually similar to a standard linear regression residual plot and can be used in the same way. From these plots, Fig. **Fig. S5 - Fig. S8**, we determine that the linearity assumption holds well, so logistic regression is a suitable model for our study.

### Supplementary material S4. Modelling method selection

We modelled the probability of initial attack success using four methods: logistic regression, balanced random forests, generalised additive models (GAMs) and mixed models. Logistic regression is the simplest and most interpretable method but is the least flexible. Diagnostic plots for the logistic regression model are given in Supplementary material S2. The rest of the models allow more flexibility and so can capture more complex relationships between the input variables and the outcome, so we compared these methods to logistic regression. In this supplementary material, we demonstrate that the more flexible models do not give much performance improvement. This result aligns with the residual plots in Supplementary material S3 which indicate that the linearity assumption for logistic regression is adequately satisfied.

#### Balanced random forest

Since the data is unbalanced (there are many more controlled fires than uncontrolled), we used the Python package imbalanced-learn which has random forest algorithms that deal with unbalanced data. We split the training data set into training (80%) and validation (20%) sets for hyperparameter tuning and chose the F1 metric for the optimisation. The F1 score is the harmonic mean of the precision and recall scores, so it is an appropriate goodness measure for models with unbalanced data.

The diagnostic plots are shown in Fig. Fig. S9 -Fig. S12. For balanced random forest models, the decision threshold is always 0.5, which is why the distributions are centred around a probability of a half. Comparing the AUC and AP with the logistic regression values in Fig. S1 - S4 indicates that the random forest models are on par with the logistic regression models.

#### Generalised additive models

We fit generalised additive models (GAMs) using R and the package mgcv (Wood 2011, 2017). The number of knots were chosen using the automatic selection of the package and then checked using gam.check. In all cases, the effective degrees of freedom were much lower than the maximal number of knots tested by the function, indicating there was no need to manually increase the number of knots tested. The model performance is similar to the performance obtained by the linear logistic regression model, see Fig. S13 - S16.

### Supplementary material S5. Detailed review of fires

In order to evaluate the model’s performance, we identified fires where the model’s prediction were unexpected compared to the actual outcome. We reviewed situation reporting and radio logs to identify any factors influencing the initial attack outcome, see **Table S13**.

## Figures

**Fig. S1.**
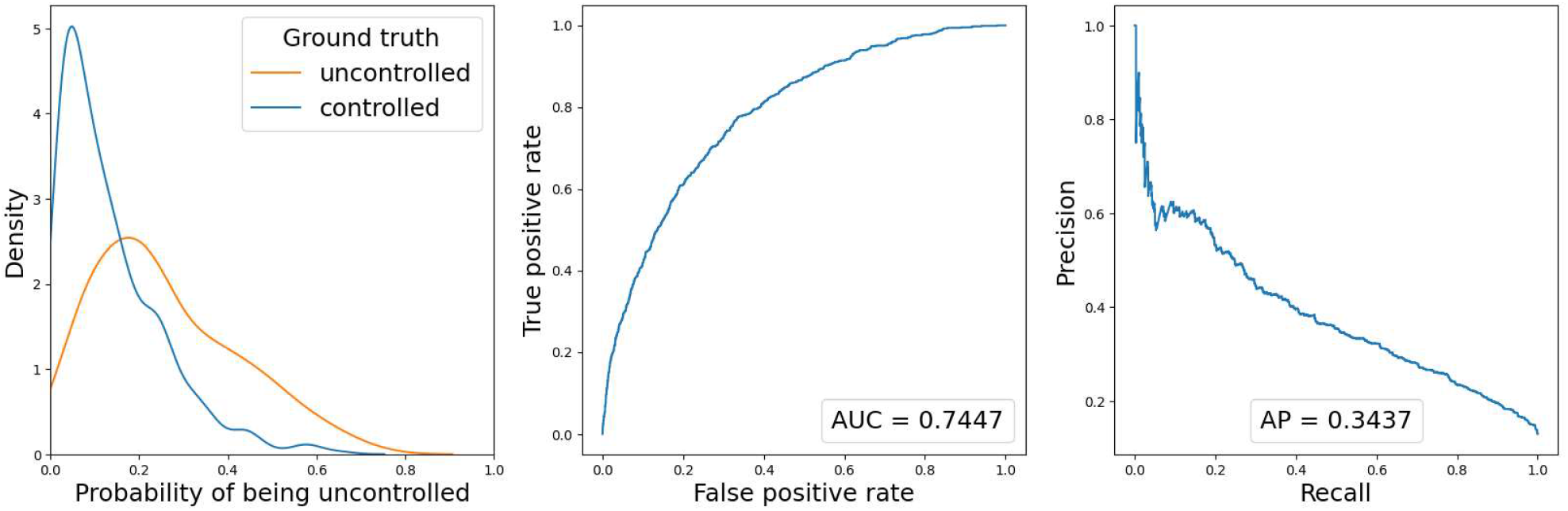
Diagnostic plots of the model trained on grass fires, estimating the probability of being uncontrolled (containment time will be greater than 2 h). The leftmost plot shows a kernel density plot of the probabilities predicted by the model for the test set, split by the true outcome. The further separated the blue and orange graphs, the better the model is at distinguishing between fires that were contained (blue) and uncontained (orange). The middle plot is a Receiver Operator Curve and the rightmost a Precision-Recall curve. Both measure model performance, but the Precision-Recall curve is considered better for unbalanced data such as the data in this study.

**Fig. S2.**
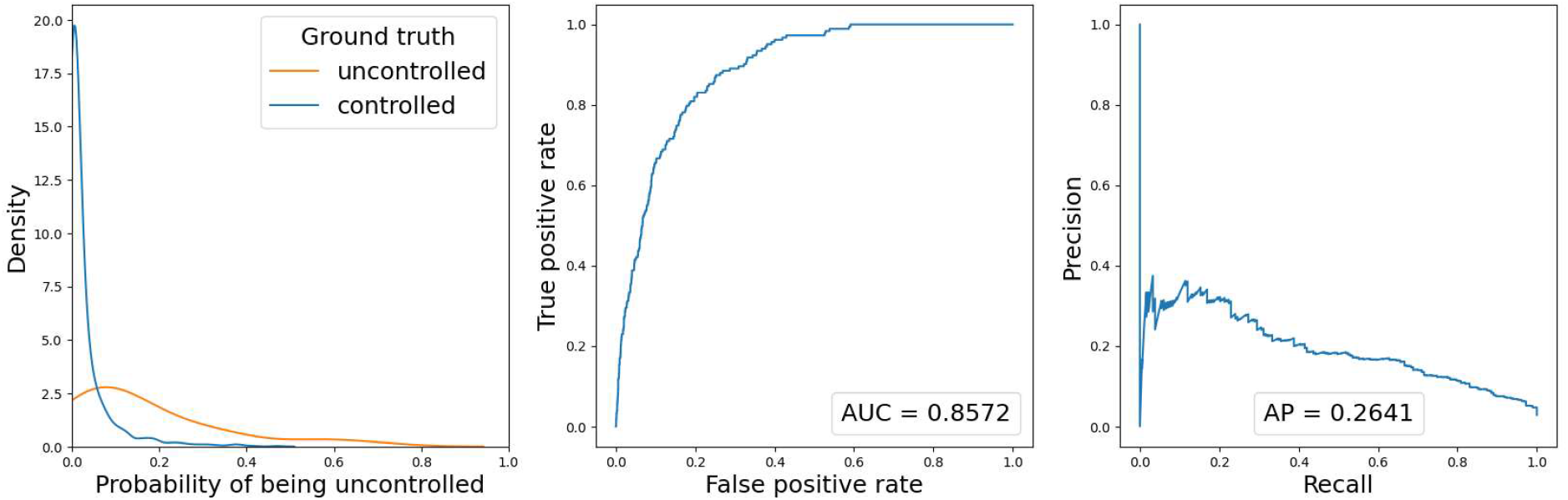
Diagnostic plots of the model trained on grass fires, estimating the probability of being uncontrolled (final fire area will be greater than 100 ha). The leftmost plot shows a kernel density plot of the probabilities predicted by the model for the test set, split by the true outcome. The further separated the blue and orange graphs, the better the model is at distinguishing between fires that were contained (blue) and uncontained (orange). The middle plot is a Receiver Operator Curve and the rightmost a Precision-Recall curve. Both measure model performance, but the Precision-Recall curve is considered better for unbalanced data such as the data in this study.

**Fig. S3.**
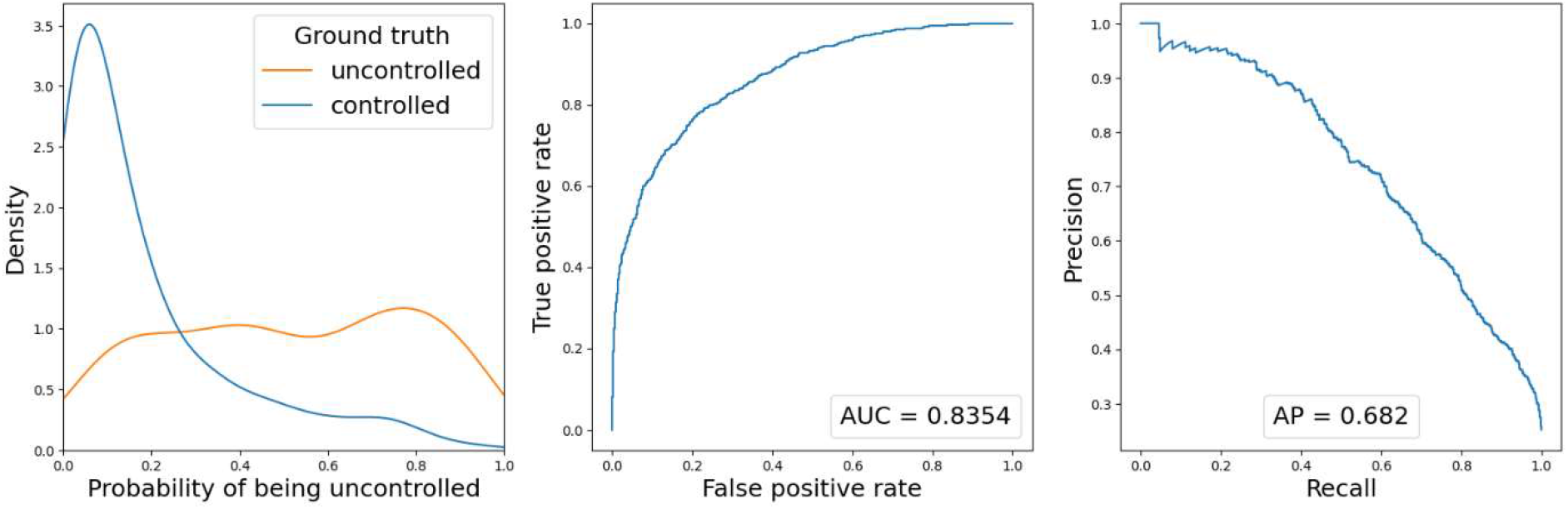
Diagnostic plots of the model trained on forest fires, estimating the probability of being uncontrolled (containment time will be greater than 4 h). The leftmost plot shows a kernel density plot of the probabilities predicted by the model for the test set, split by the true outcome. The further separated the blue and orange graphs, the better the model is at distinguishing between fires that were contained (blue) and uncontained (orange). The middle plot is a Receiver Operator Curve and the rightmost a Precision-Recall curve. Both measure model performance, but the Precision-Recall curve is considered better for unbalanced data such as the data in this study.

**Fig. S4.**
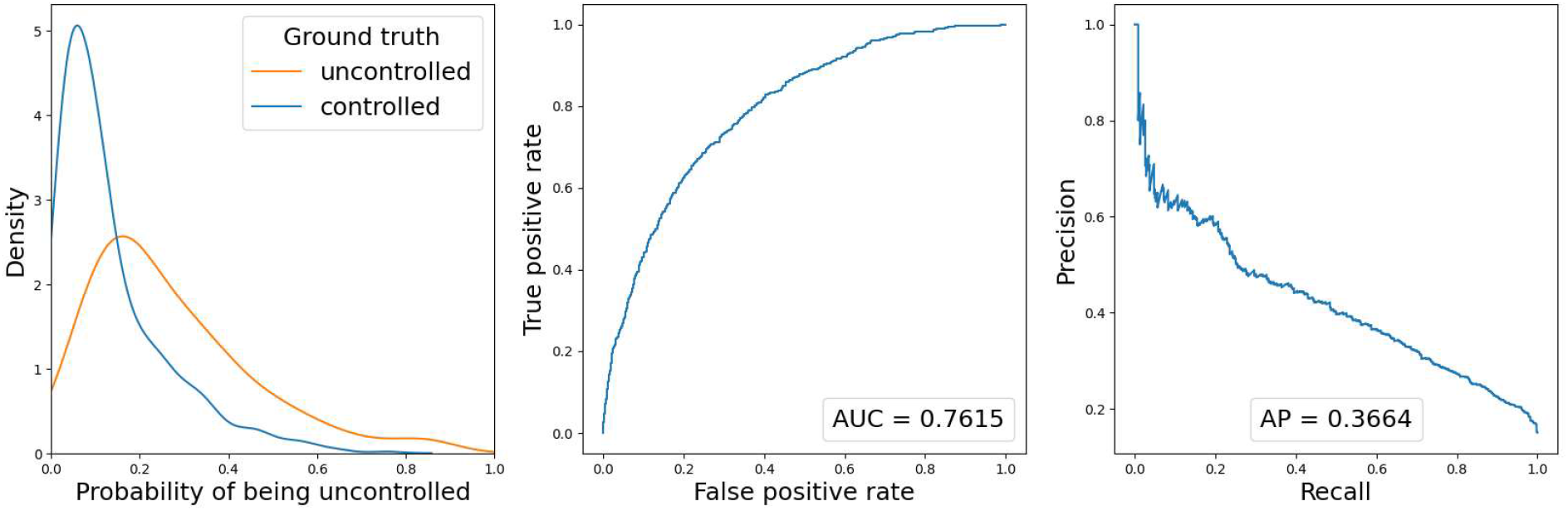
Diagnostic plots of the model trained on forest fires, estimating the probability of being uncontrolled (final fire area will be greater than 5 ha). The leftmost plot shows a kernel density plot of the probabilities predicted by the model for the test set, split by the true outcome. The further separated the blue and orange graphs, the better the model is at distinguishing between fires that were contained (blue) and uncontained (orange). The middle plot is a Receiver Operator Curve and the rightmost a Precision-Recall curve. Both measure model performance, but the Precision-Recall curve is considered better for unbalanced data such as the data in this study.

**Fig. S5.**
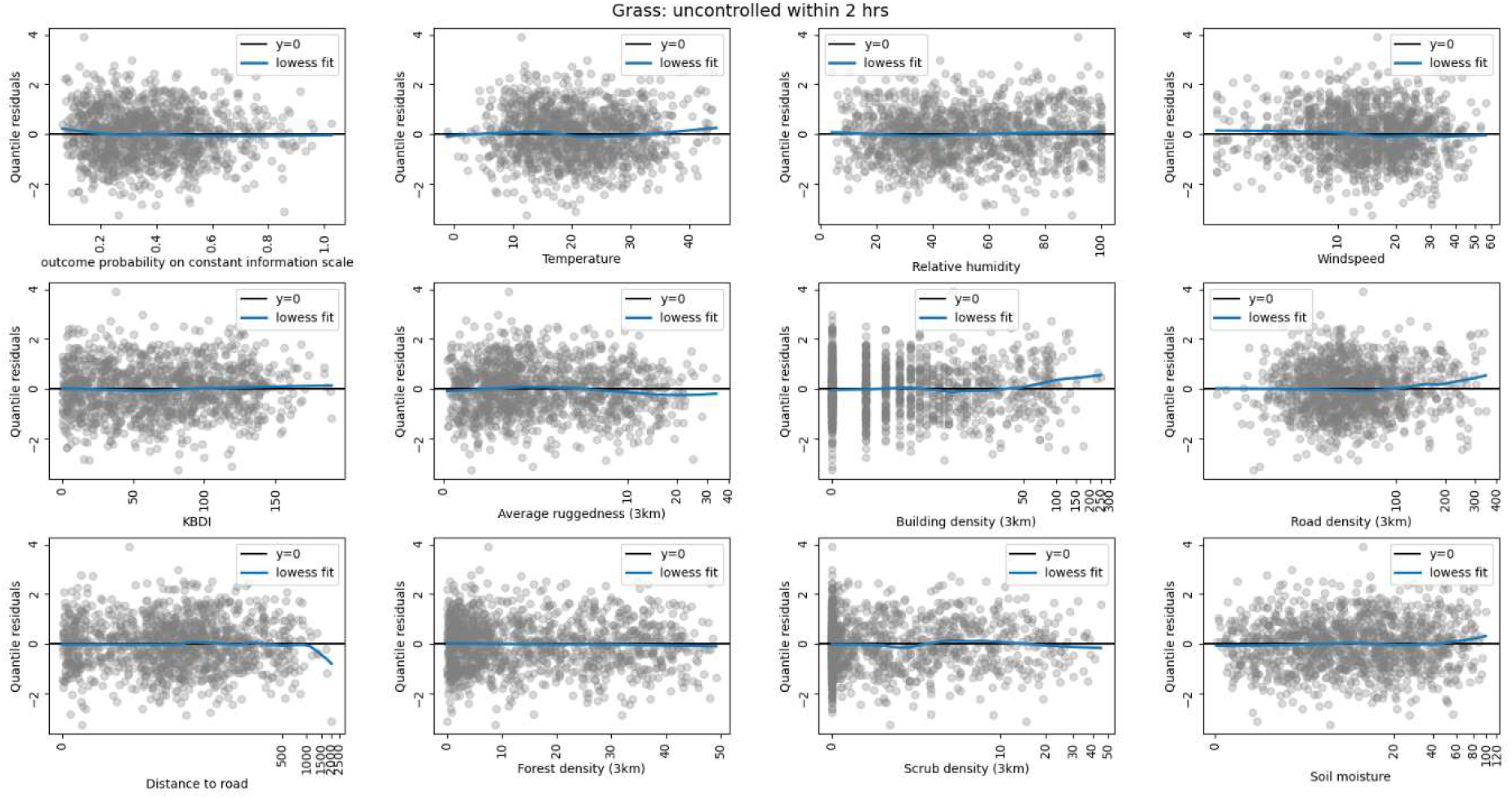
Randomised quantile residual plots for the model trained on grass fires, estimating the probability of being uncontrolled (containment time will be greater than 2 h). These plots show that the linearity assumption holds well for the variables in our model because the residuals are scattered evenly above and below the horizontal axis.

**Fig. S6.**
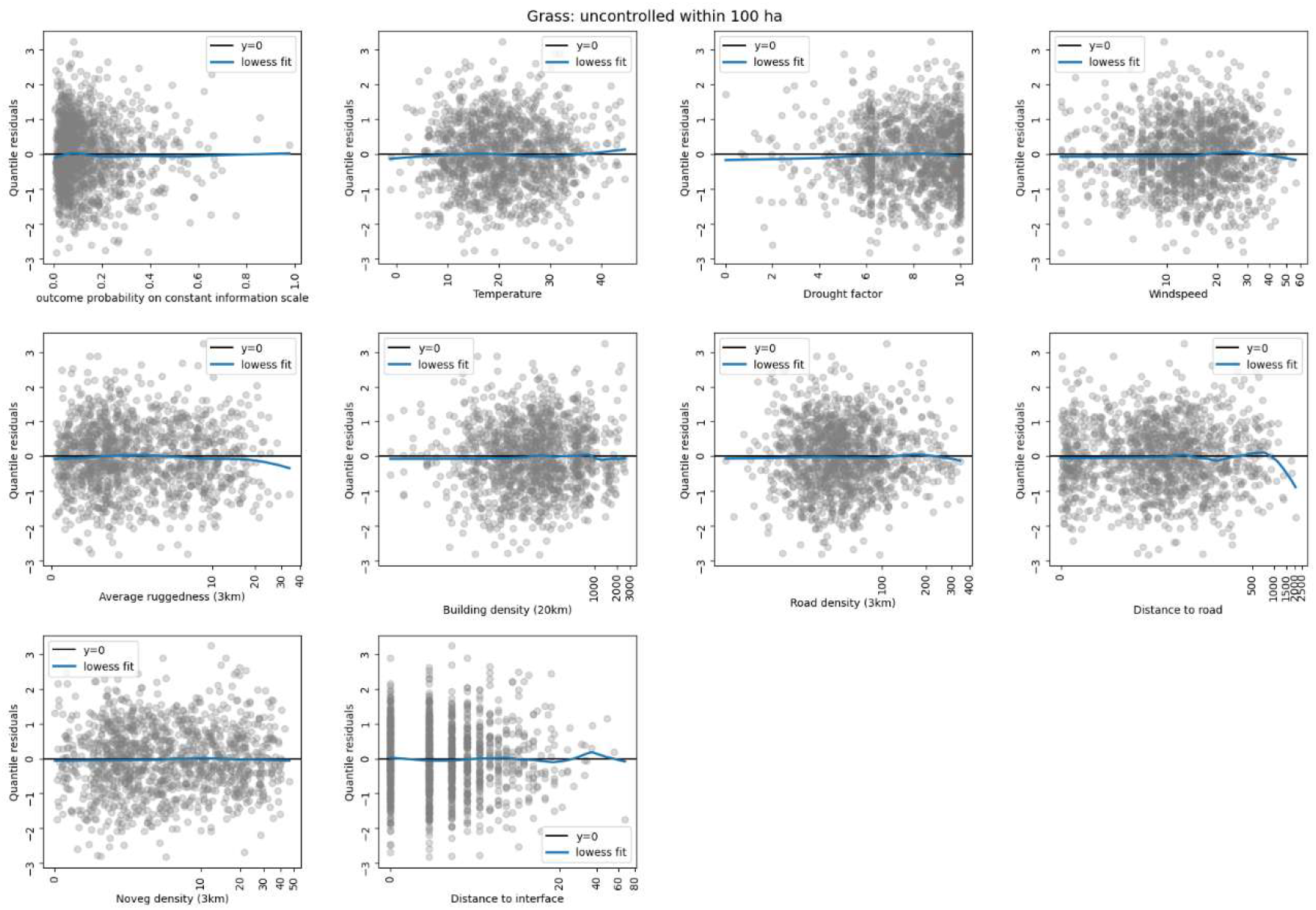
Randomised quantile residual plots for the model trained on grass fires, estimating the probability of being uncontrolled (final fire area will be greater than 100 ha). These plots show that the linearity assumption holds well for the variables in our model because the residuals are scattered evenly above and below the horizontal axis.

**Fig. S7.**
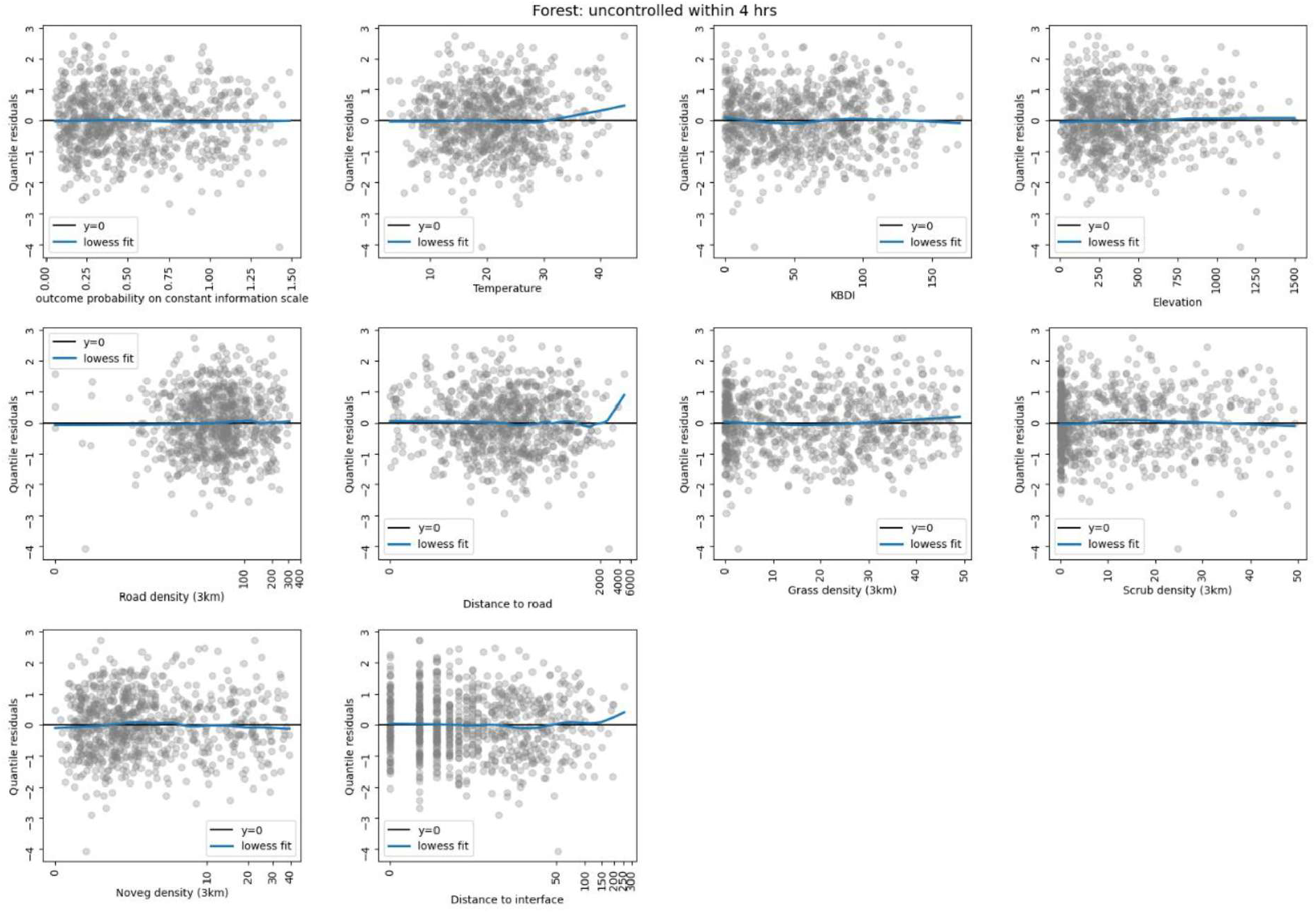
Randomised quantile residual plots for the model trained on forest fires, estimating the probability of being uncontrolled (containment time will be greater than 4 h). These plots show that the linearity assumption holds well for the variables in our model because the residuals are scattered evenly above and below the horizontal axis.

**Fig. S8.**
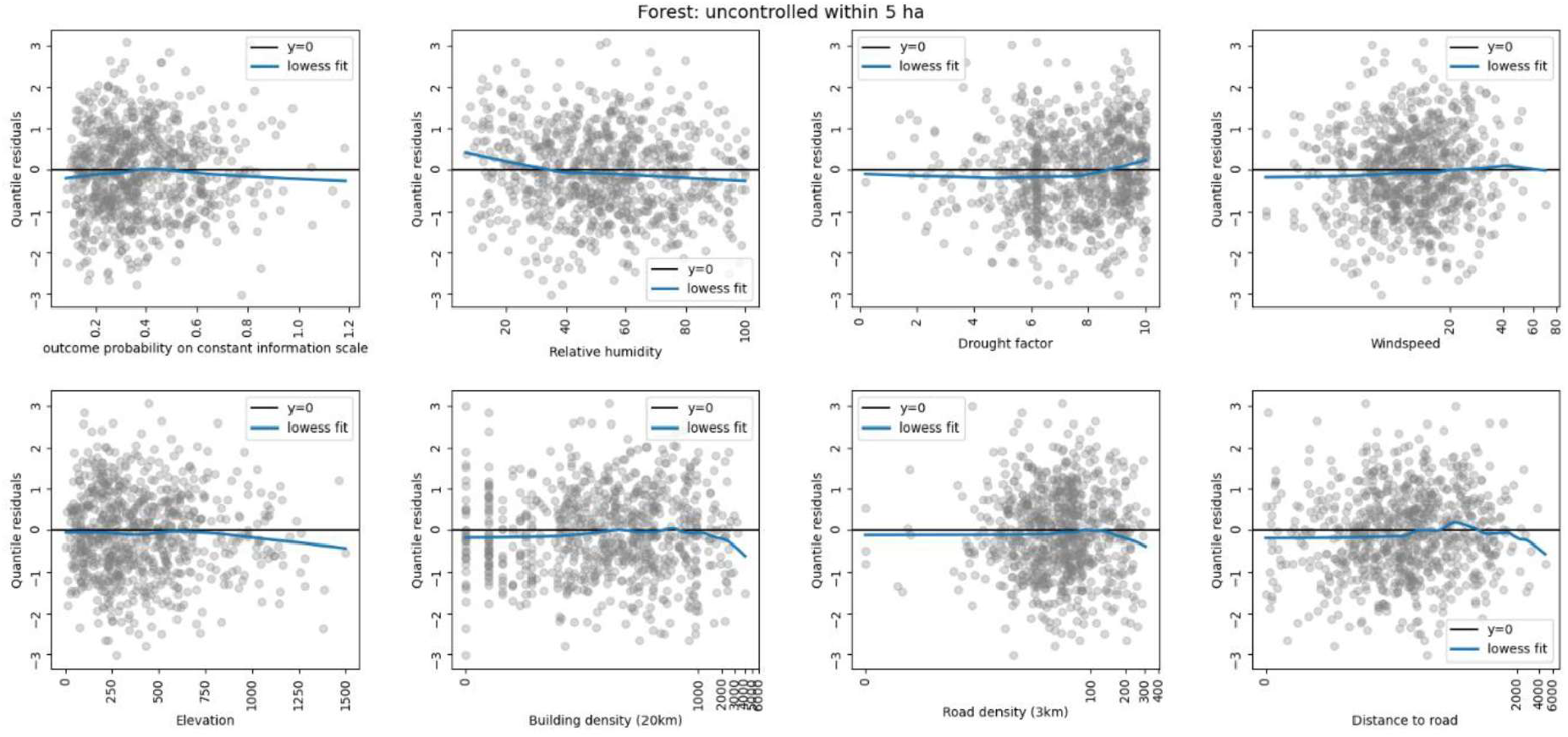
Randomised quantile residual plots for the model trained on forest fires, estimating the probability of being uncontrolled (final fire area will be greater than 5 ha). These plots show that the linearity assumption holds well for the variables in our model because the residuals are scattered evenly above and below the horizontal axis.

**Fig. S9.**
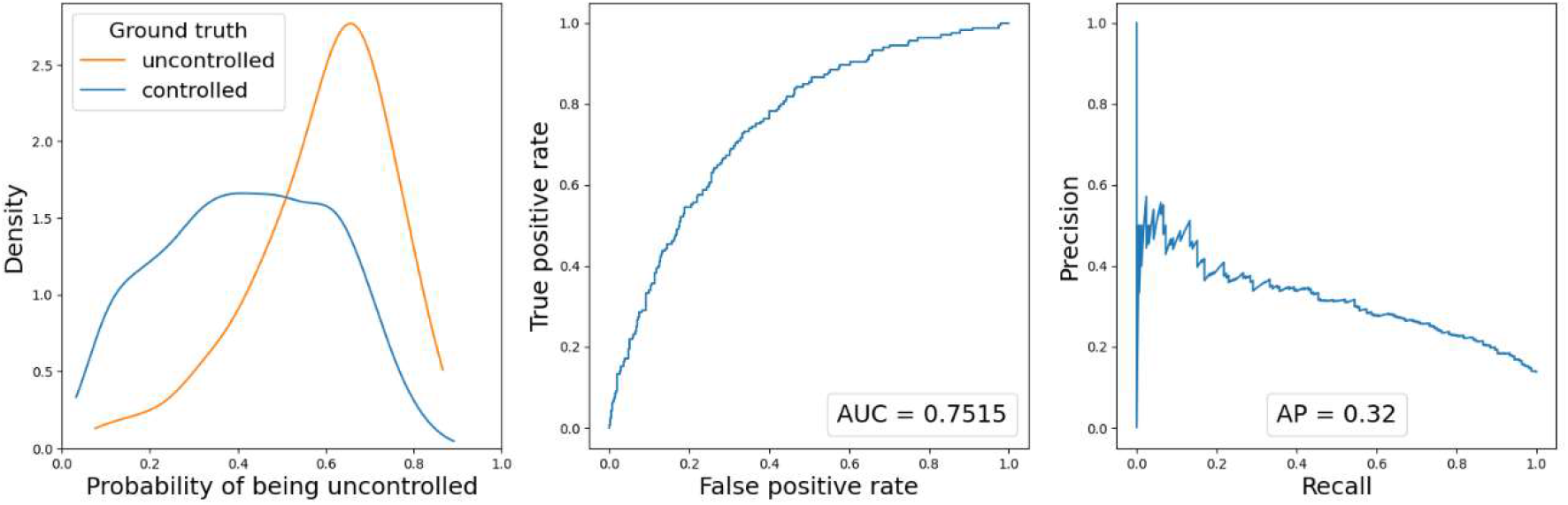
Diagnostic plots for the balanced random forest model trained on grass fires, estimating the probability of being uncontrolled (containment time will be greater than 2 h).

**Fig. S10.**
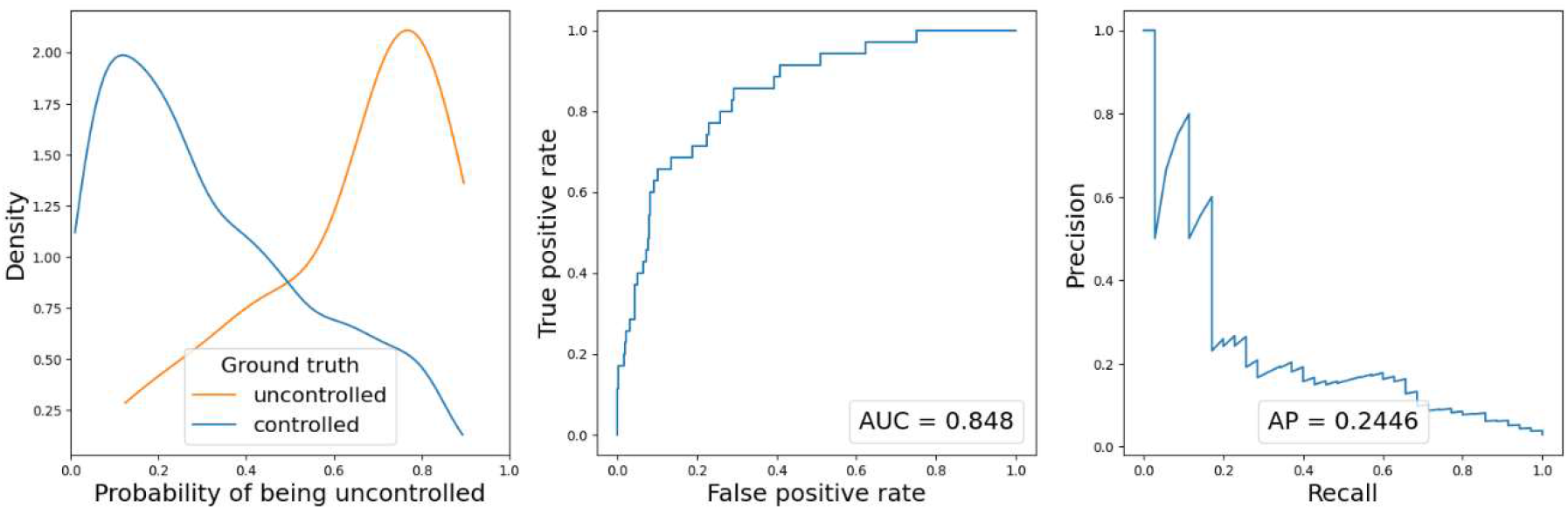
Diagnostic plots for the balanced random forest model trained on grass fires, estimating the probability of being uncontrolled (final fire area will be greater than 100 ha).

**Fig. S11.**
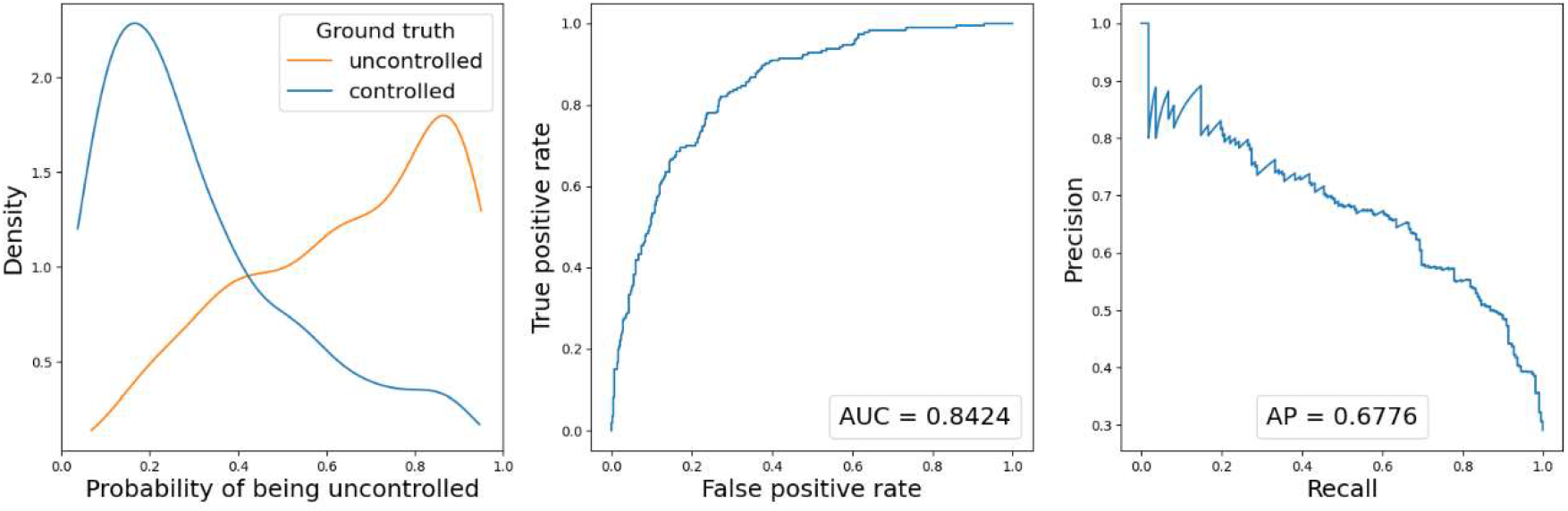
Diagnostic plots for the balanced random forest model trained on forest fires, estimating the probability of being uncontrolled (containment time will be greater than 4 h).

**Fig. S12.**
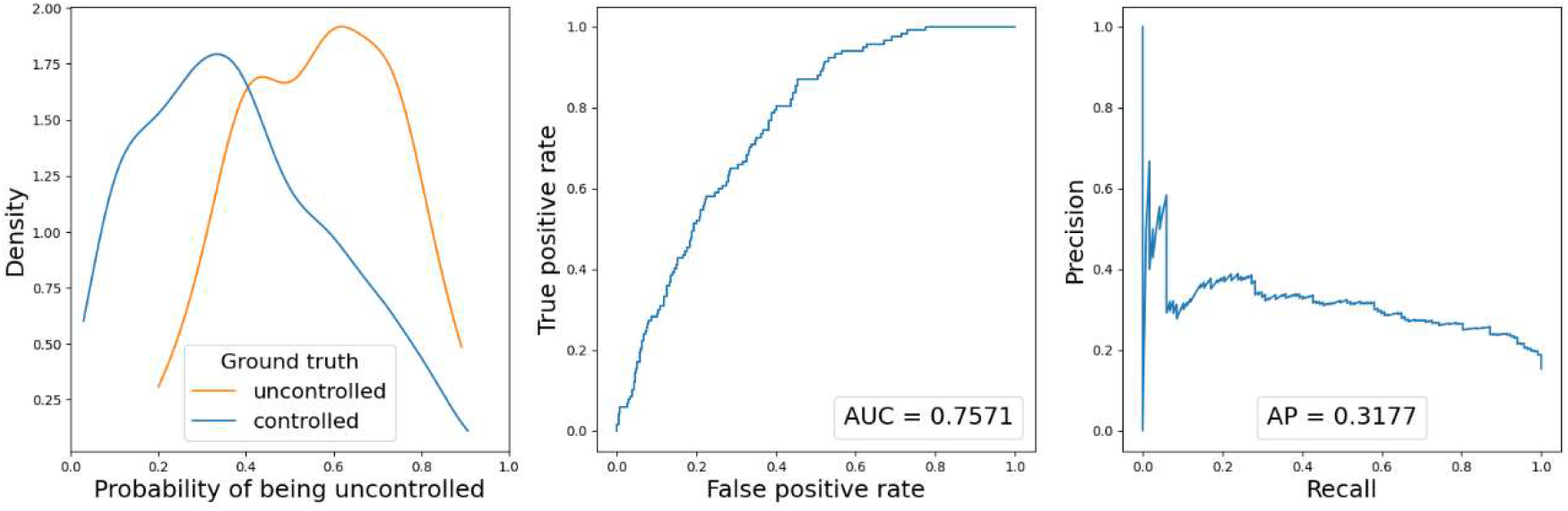
Diagnostic plots for the balanced random forest model trained on grass fires, estimating the probability of being uncontrolled (final fire area will be greater than 5 ha).

**Fig. 13.**
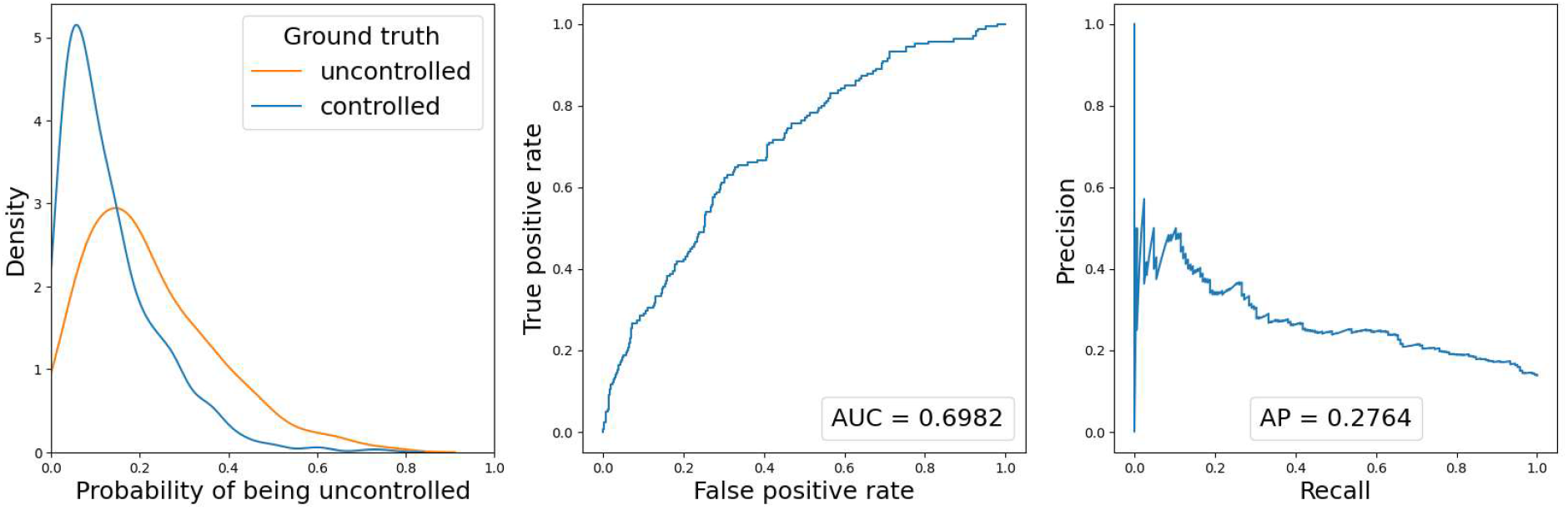
Diagnostic plots for the GAM trained on grass fires, estimating the probability of being uncontrolled (containment time will be greater than 2 h).

**Fig. 14.**
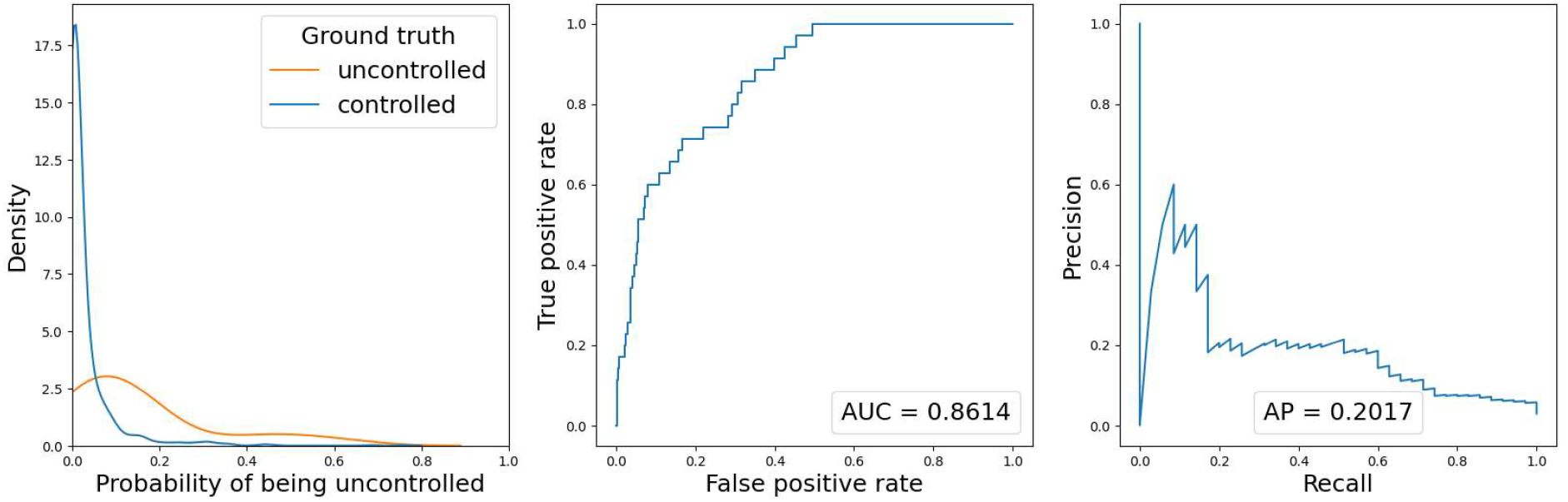
Diagnostic plots for the GAM trained on grass fires, estimating the probability uncontrolled (final fire area will be greater than 100 ha).

**Fig. 15.**
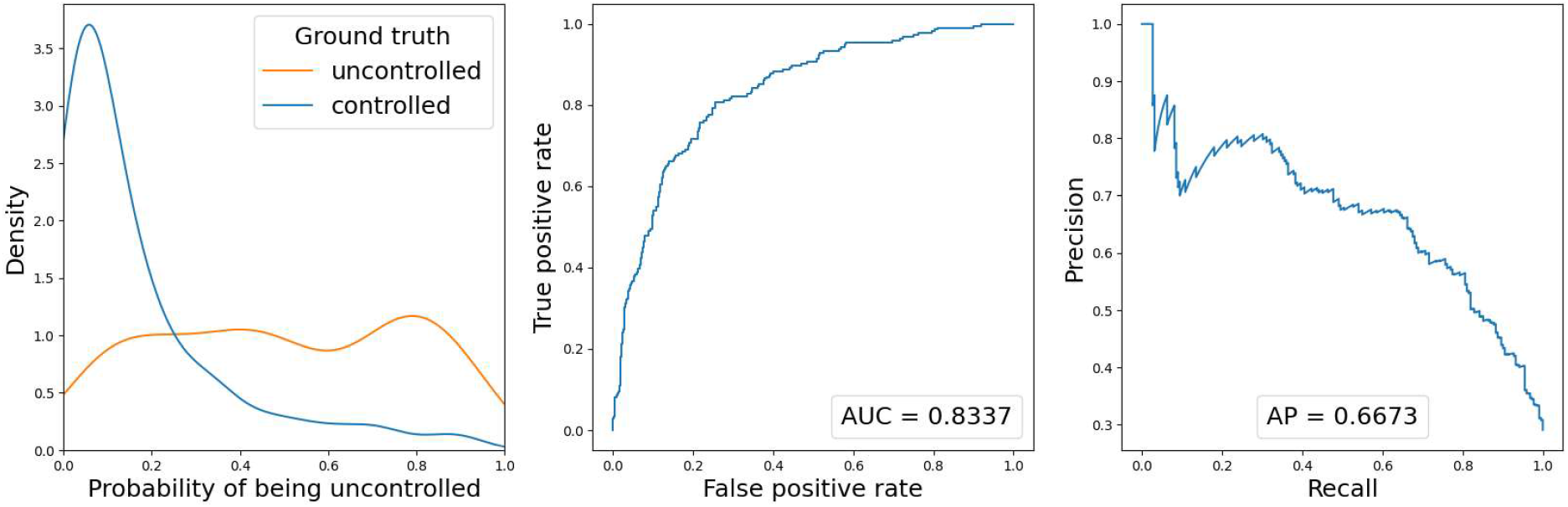
Diagnostic plots for the GAM trained on forest fires, estimating the probability uncontrolled (containment time will be greater than 4 h).

**Fig. 16.**
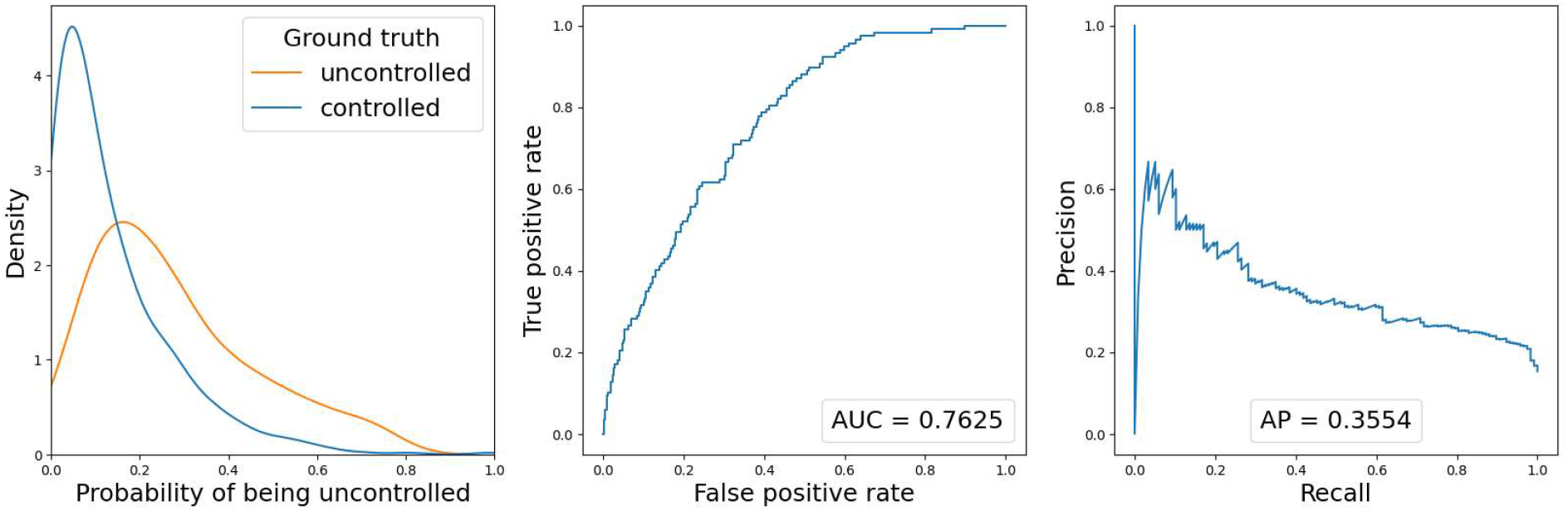
Diagnostic for the GAM trained on forest fires, estimating the probability uncontrolled (final fire area will be greater than 5 ha).

## Tables

**Table S11.** Test train split by fire season.

| Split | Season | Season with major fires |
| --- | --- | --- |
| Train (76%) | 2009-10 |  |
|  | 2010-11 |  |
|  | 2011-12 |  |
|  | 2012-13 |  |
|  | 2013-14 |  |
|  | 2014-15 |  |
|  | 2015-16 |  |
|  | 2018-19 |  |
|  | 2019-20 | ✓ |
|  | 2020-21 |  |
|  | 2021-22 |  |
|  | 2022-23 |  |
|  | 2023-24 |  |
| Test (24%) | 2007-08 |  |
|  | 2008-09 | ✓ |
|  | 2017-18 |  |
|  | 2020-21 |  |

**Table S12.** Percentage of fires (rows of data) in the train and test sets for each model. Each row sums to 100%.

| Fuel | Train | Test |
| --- | --- | --- |
| Grass | 84% | 16% |
| Forest | 80% | 20% |

**Table S13.**
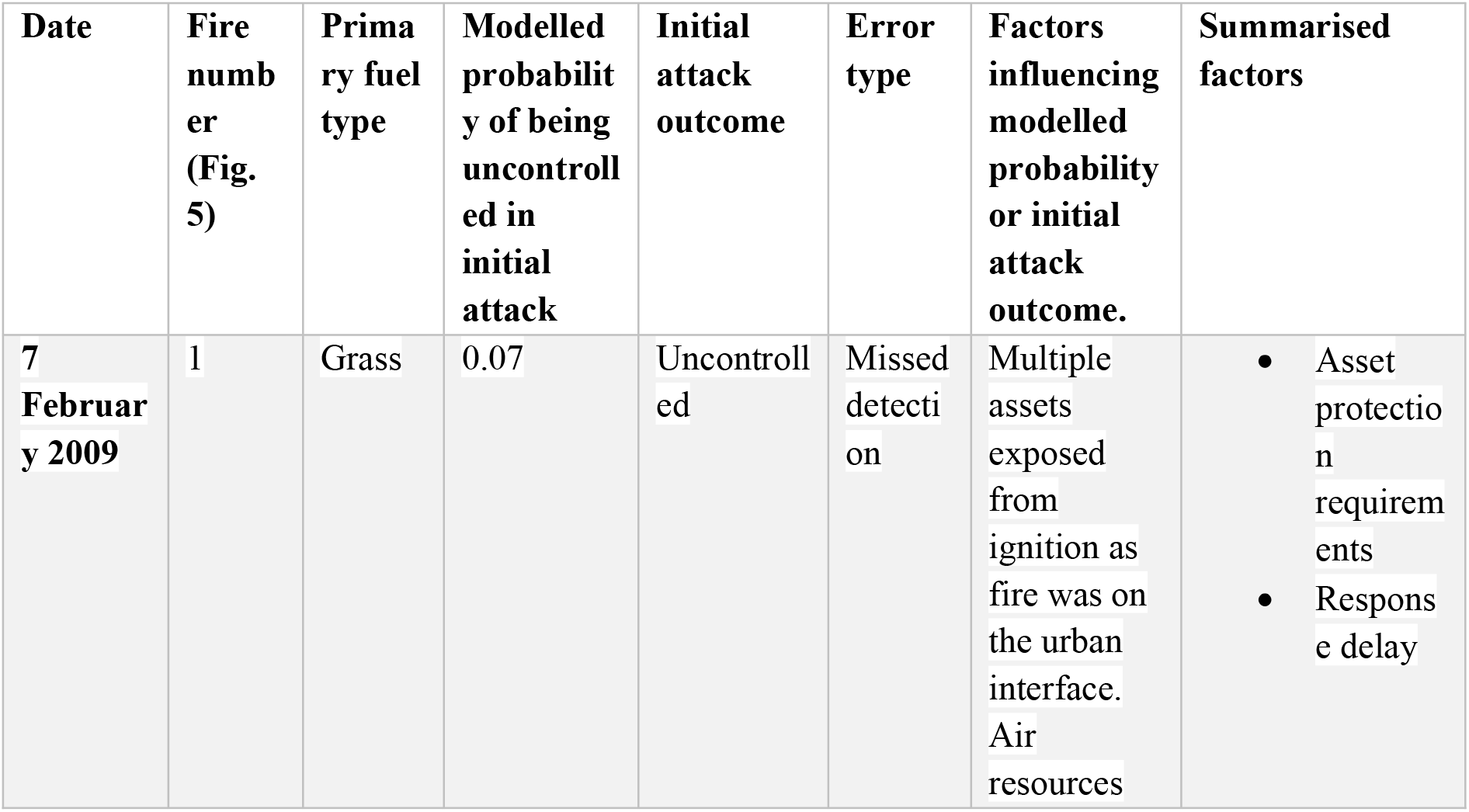
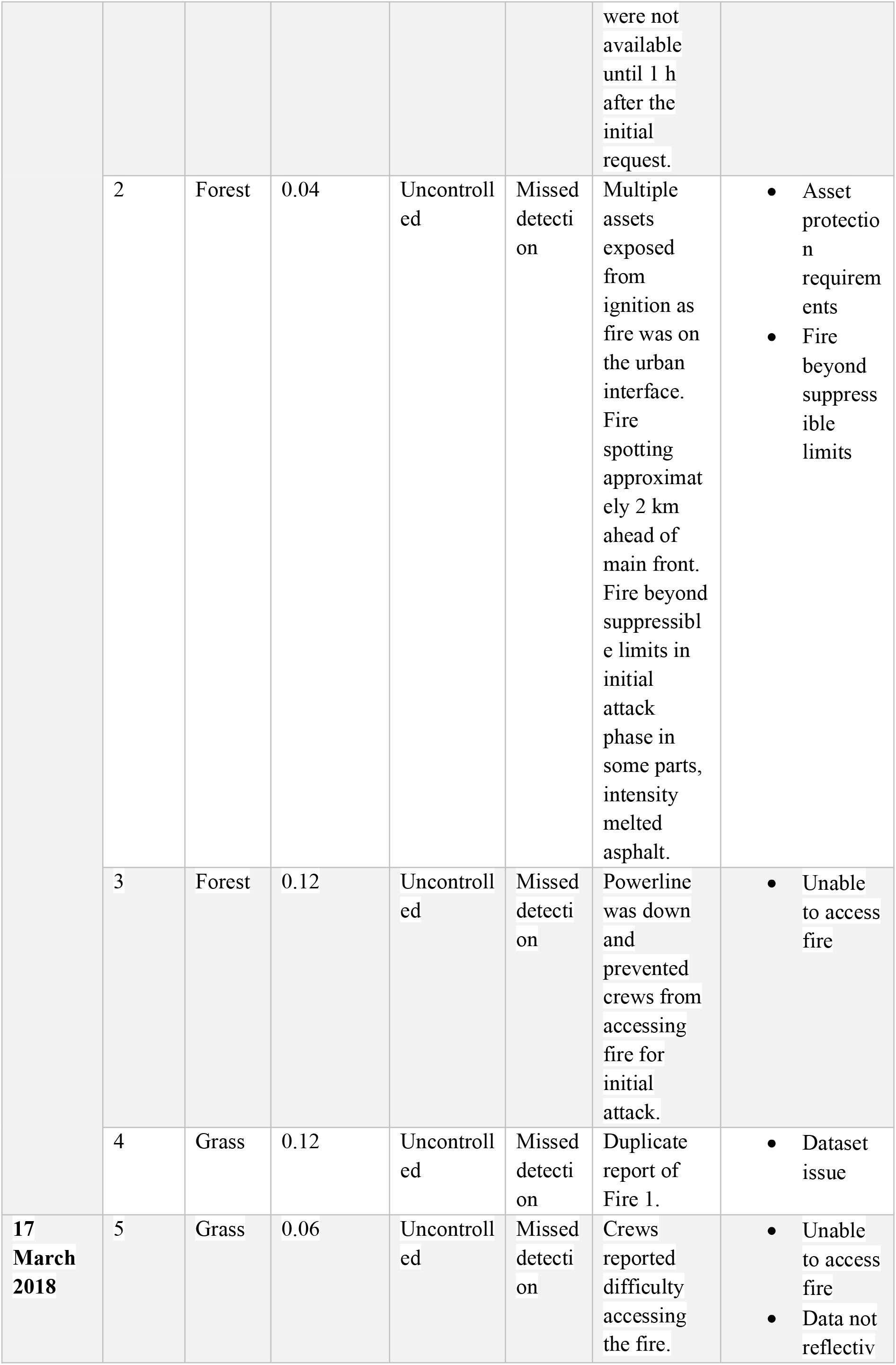
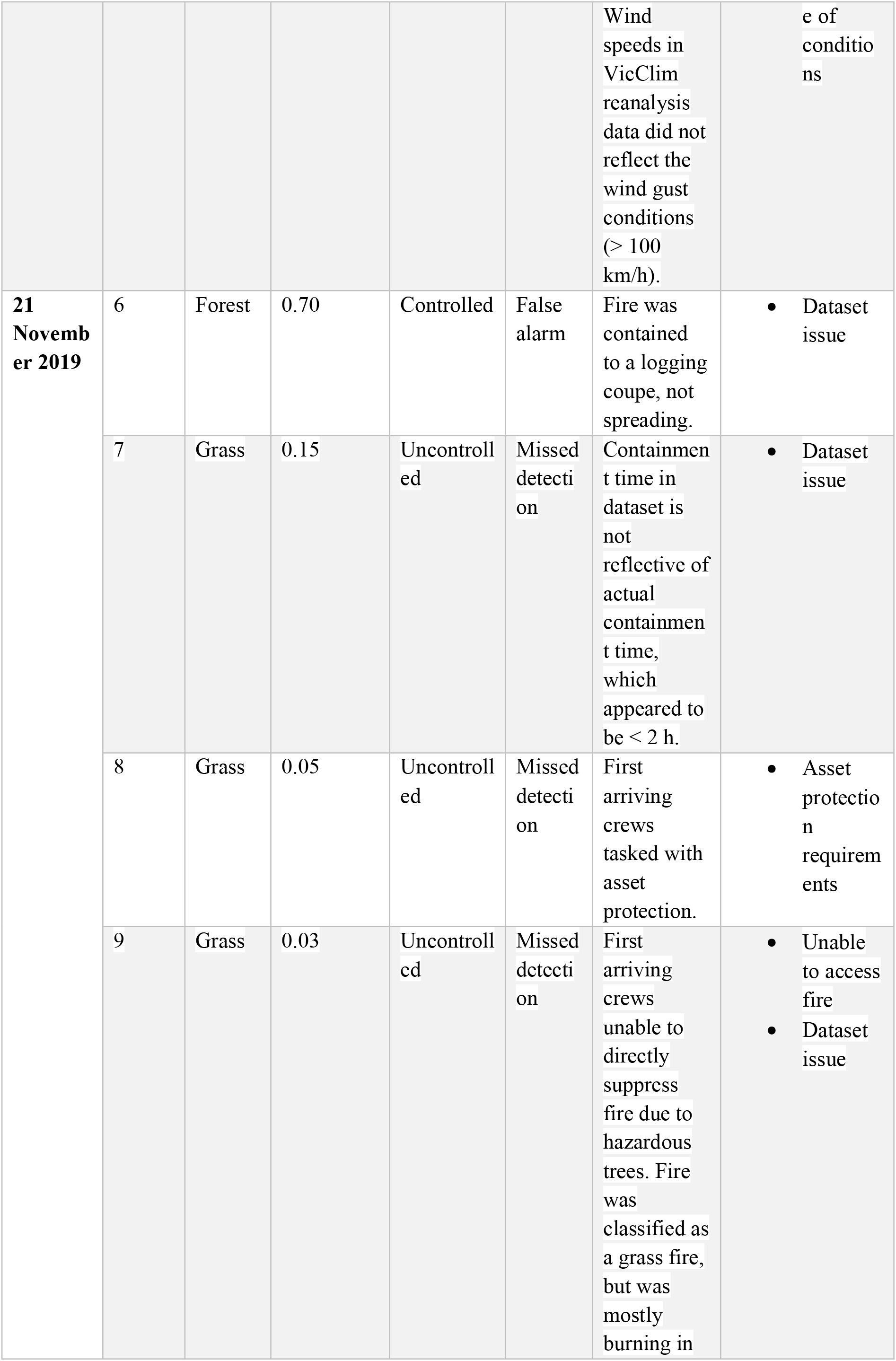
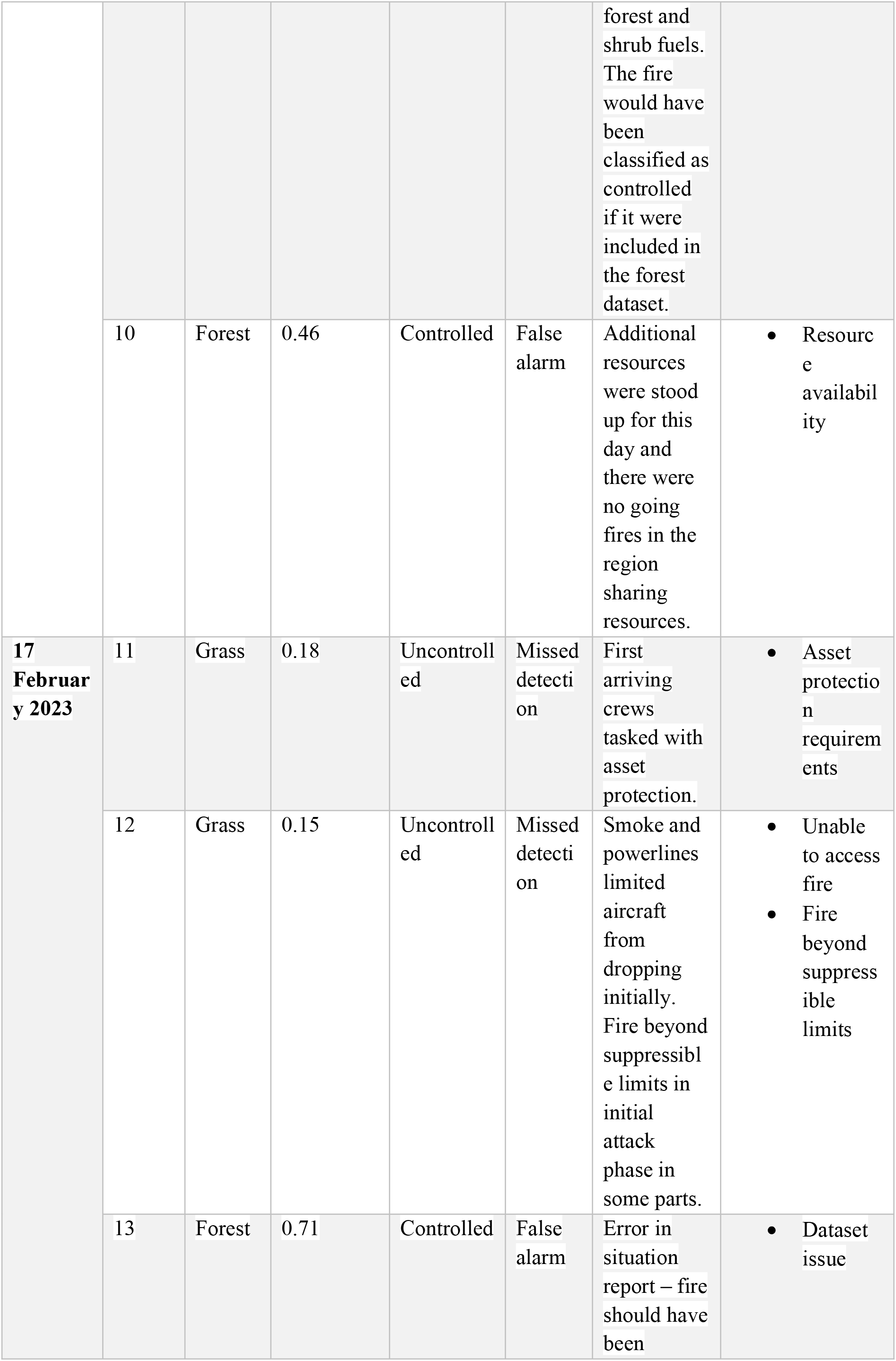
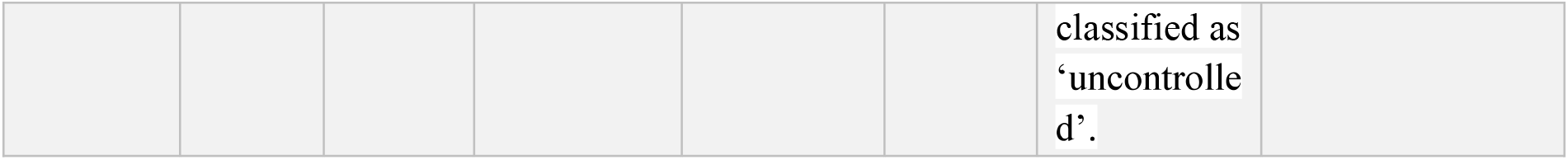
Detailed review of fires with unexpected modelled predictions relative to the initial attack outcome.

